# Axin proteolysis by Iduna is required for the regulation of stem cell proliferation and intestinal homeostasis in *Drosophila*

**DOI:** 10.1101/296830

**Authors:** Yetis Gultekin, Hermann Steller

## Abstract

The self-renewal of intestinal stem cell is controlled by Wingless/Wnt-β catenin signaling both in *Drosophila* and mammals. Since Axin is a rate-limiting factor in Wingless signaling its regulation is essential. Iduna is an evolutionarily conserved ubiquitin E3 ligase that has been identified as a critical regulator for degradation of ADP-ribosylated Axin and thus of Wnt/β-catenin signaling. However, its physiological significance remains to be demonstrated. Here, we generated loss-of-function mutants of *Iduna* to investigate its physiological role in *Drosophila*. Genetic depletion of *Iduna* causes the accumulation of both Tankyrase and Axin. Increase of Axin protein in enterocytes non-autonomously enhanced stem cell divisions in the *Drosophila* midgut. Enterocytes secreted Unpaired and thereby stimulated the activity of the JAK-STAT pathway in intestinal stem cells. A decrease in *Axin* gene expression suppressed both the over-proliferation of stem cells and restored their numbers to normal levels in *Iduna* mutants. These findings suggest that Iduna-mediated regulation of Axin proteolysis is essential to maintain tissue homeostasis in the *Drosophila* midgut.

## Introduction

The evolutionarily conserved Wnt/β-catenin signaling pathway is a main regulator of animal development. It controls proliferation, differentiation and regeneration of adult tissues (Herr et al., 2012; Nusse and Clevers, 2017). The Wingless pathway is also involved in adult tissue self-renewal in *Drosophila* (Lin et al., 2008). Genetic depletion of proteins in the Wingless pathway, such as *Tcf, arr, dsh* and *pygo,* leads to inhibition of Wingless signaling activation which in turn causes over-proliferation of stem cells in the *Drosophila* midgut (Kramp et al., 2002; Wang et al., 2016a and 2016b; Tian et al., 2016). However, inactivation of Wnt signaling in the small intestine of mice decreases the proliferative potential of stem cells (Fevr et al., 2007; Korinek et al., 1998). On the other hand, mutations resulting in the over-activation of the Wnt/β-catenin pathway promote tumorigenesis (Clevers and Russe, 2012; Andreu et al., 2005; Korinek et al., 1997 and 1998; Morin et al., 1997). For instance, mutations on *adenomatous polyposis coli (APC)* gene cause a hereditary colorectal cancer syndrome called familial adenomatous polyposis (Kinzler et al., 1991; Nishisho et al., 1991). Axin loss-of-function mutations are found in hepatocellular carcinomas, while oncogenic β-catenin mutations are described in colon cancer and melanoma (Rubinfeld et al., 1997). Consequently, intense efforts have been made to target this pathway for therapeutic purposes (Clevers and Russe, 2012).

A key feature of the Wnt/β-catenin pathway is the regulated proteolysis of the downstream effector β-catenin by the β-catenin degradation complex. The principal components of this complex are adenomatous polyposis coli (APC), Axin and Glycogen synthase kinase 3β (GSK3β) (Kramps et al., 2002; Hamada et al., 1999; Salic et al., 2000; Lee at al., 2003). Axin, a critical scaffold protein in the β-catenin degradation complex, is the rate-limiting factor of Wnt signaling and its protein levels are regulated by the Ubiquitin-Proteasome System (UPS) (Li et al., 2012). Axin is targeted for degradation by the combined action of the poly-ADP-ribose polymerase Tankyrase (TNKS) and the ubiquitin E3-ligase Iduna/Ring finger protein 146 (RNF146) (Zhang et al., 2011). Both genetic and pharmacological studies suggest that UPS-dependent degradation of Axin occurs in a specific temporal order. Iduna initially exists in an inactive state, but binding to its iso-or poly-ADP-ribosylated targets causes allosteric activation of the enzyme (DaRosa et al., 2015). In the first step, TNKS binds to Axin and ADP-ribosylates Axin using NAD^+^. Then, Iduna recognizes and binds to ADP-ribosylated Axin via its WWE domain and poly-ubiquitylates Axin. Following the ADP-ribosylation and ubiquitination, post-translationally modified Axin is rapidly degraded by the proteasome (DaRosa et al., 2015; Wang et al., 2016a and 2016b; Croy et al., 2016; Callow et al., 2011). This tight control suggests an important function for Iduna to regulate the Wnt-β catenin pathway.

Since the stability of Axin is partially regulated by TNKS-mediated ADP ribosylation, specific small-molecule inhibitors have been developed to inhibit Wnt-signaling (Lu et al., 2009; Huang et al., 2009). For example, XAV939 targets the ADP-ribose polymerase activity of TNKS and increases Axin levels, which in turn destabilizes β-catenin to inhibit Wnt signaling (Huang et al., 2009). There are two TNKS isoforms in mammalian cells (Hsiao et al., 2006). *Tnks1*^*-/-*^ and *Tnks2*^*-/-*^ mice are overall normal; however, double knock-out of *Tnks*^*1/2*^ causes early embryonic lethality, which indicates their redundancy in mouse development (Hsiao et al., 2006; Chiang et al., 2008). On the other hand, inactivation of the single *Drosophila Tnks* gene produces viable flies that have slightly increased Axin levels and abnormal proliferation of intestinal stem cells, but otherwise display no overt defects (Wang et al., 2016a and 2016b; Feng et al., 2014; Yang et al., 2016; Tian et al., 2016). On the other hand, the exact physiological function of Iduna remains to be determined. In order to address this question, we generated and characterized *Drosophila* Iduna loss-of-function mutants and demonstrate a critical function of this pathway for stem cells in the *Drosophila* intestinal tract.

The *Drosophila* genomes encode four isoforms of *CG8786/Iduna/RNF146*, which is evolutionarily conserved from *Drosophila* to human. In this study, we concentrated on the physiological function of Iduna in the adult *Drosophila* midgut, which shares several striking similarities with the mammalian small intestine but offers greater anatomical and genetic accessibility (Micchelli et al., 2005; Ohlstein et al., 2005; Markstein et al., 2013). Under normal conditions, Wingless signaling controls stem cell proliferation and cell fate specification in adult midgut (Tian et al., 2016). Here, we show that Iduna has a physiological function to regulate the proteolysis of both TNKS and Axin. Inactivation of *Iduna* results in increased numbers of midgut stem cells and progenitors due to over-proliferation. We find that Axin accumulation in enterocytes promotes the secretion of Unpaired, a cytokine that binds to the Domeless receptor and activates the JAK-STAT pathway in stem cells and thereby promotes stem cell division. Significantly, reducing *Axin* expression by half restores the numbers of ISC. These findings indicate that regulation of Axin proteolysis by Iduna is necessary to control intestinal homeostasis in *Drosophila,* and it provides physiological evidence for the idea that the function of Tnks and Iduna/RNF146 is tightly coupled.

## Results

### Iduna plays a role in Axin degradation

To examine the *in vivo* function of *Drosophila* Iduna, CRISPR-Cas9 genome editing was used to generate *Iduna* mutants. *Iduna* is located on the 3^rd^ chromosome of *Drosophila*. We designed a specific guide RNA that targets *Iduna*‵s first exon and identified two mutant alleles by Sanger sequencing: *Iduna*^*17*^ and *Iduna*^*78*^, which have 4-nucleotides and 2-nucleotides deletions, respectively (Fig 1A). These deletions are close to the translation start side of *Iduna*. Next we assessed the levels of mRNA and protein expression in these mutants. Using reverse transcript PCR analysis, we found significantly reduced amounts of *Iduna* transcripts in the *Iduna*^*78*^ mutant. On the other hand, we were unable to detect any *Iduna B* and *C/G* transcripts in the *Iduna*^*17*^ allele (Fig S1A). Moreover, no Iduna protein was detected in either of these mutants, indicating that they represent Null-mutations (Fig 1B). Finally, genetic analyses of these alleles in trans to a larger deletion (see below) indicate that both alleles are complete loss-of-function mutations. *Iduna* mutants were crossed to *Drosophila* deficiency lines [Df(3L) Exel6135, Df(3L) ED228)] and also to each other and all combinations were viable as trans-heterozygotes.

**Figure 1:**
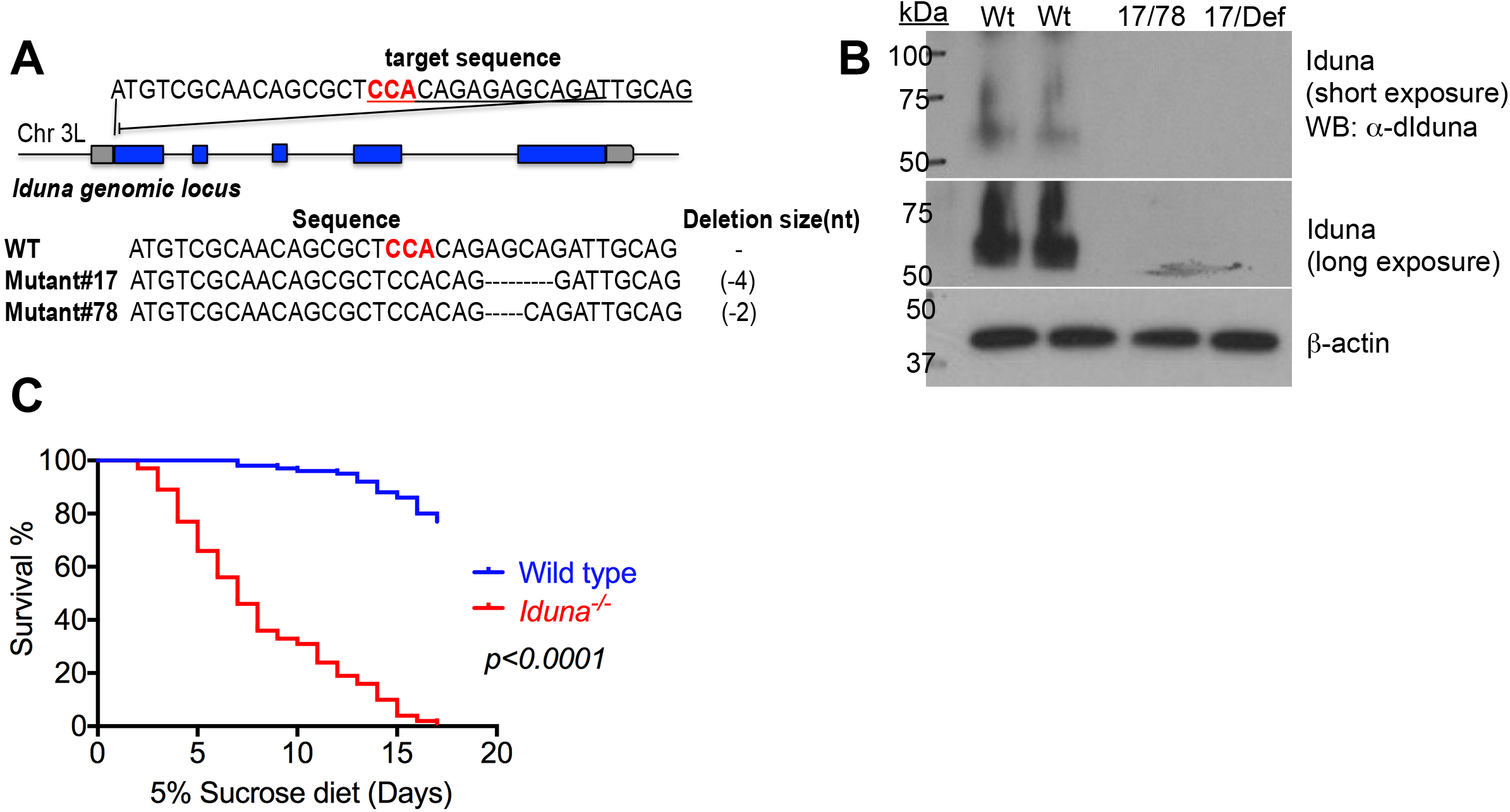
Loss-of-function mutants of *dIduna* are viable and increase TNKS and Axin protein levels. **A-** *Iduna*^*17*^ and *Iduna*^*78*^ have deletions of 4 and 2 nucleotides, respectively, introduced early stop codons and led to truncations of Iduna protein. Scheme for generation of *Iduna* loss-of-function mutants via CRISPR-Cas9 genome editing in *Drosophila.* sgRNA against Iduna was designed to generate small nucleotide deletions, close to its translation initiation site. The three nucleotides were highlighted with red to indicate the location of Cas9 cleavage site. *Iduna* loss-of-function mutants, *Iduna*^*17*^ and *Iduna*^*78*^, were isolated by Sanger sequencing**. B-** *Iduna*^*17*^ and *Iduna*^*78*^ have no detectable protein. Endogenous Iduna protein was detected by immunoblotting. Anti-Iduna antibody was generated in guinea pigs and 20µg total protein lysates of seven day-old adult females were analyzed by immunoblotting. β-actin was used as a loading control. Both *Iduna* alleles have no detectable protein and behave genetically as Null-alleles. **C-** *Iduna* mutants display increased mortality under reduced nutrient conditions. Two-day old mutant or wild type female flies were collected and kept on 5% sucrose diet at 28°C. n=100 from each genotype. **D**-There is an approximately twofold increase in the numbers of *esg>GFP* expressing stem cells/progenitors in the midgut of *Iduna* mutants compared to controls. **E-** Quantification of *esg>GFP* positive stem cells and progenitors from adult flies of indicated genotypes. n=6 from each genotype. **F**-*Iduna* inactivation increases the numbers of Arm^+^/Pros^-^ stem cells in the midguts upon nutrient deprivation. Nine day-old female flies, fed with 5% sucrose diet for seven days at 28°C, were examined in D-E and F. Posterior midguts were analyzed in this study. *p<0.001* is indicated as ***.

We examined the larval development of *Iduna* mutants and Oregon R but did not observe any differences in the numbers of hatched eggs (Fig S1B-C), pupated larvae and enclosed adult *Drosophila* (Fig S1D) between *Iduna* mutants and wild type. *Iduna*-Null adult flies had no overt morphological defects when compared with wild type controls. However they displayed increased mortality upon nutrient deprivation. We challenged mutant and wild type adult females with a 5% sucrose diet at 28°C. Two-day old adult females were placed on 5% sucrose diet at 28°C. Mutant flies died within 17 days, while 70-80% of wild type flies were still viable at this time (Fig 1C).

Iduna is one of the key components of the machinery that degrades Axin whose ADP-ribosylation by TNKS is important for mammalian Wnt-β catenin signaling (Li at al., 2012). We detected increased levels of endogenous Axin in the lysates from *Iduna* mutants midguts compared to controls (Fig 2A). Mammalian Iduna recognizes both ADP-ribosylated (ADPR) TNKS and Axin via the R163 residue in its WWE domain (Zhang et al., 2011). The R163 residue is conserved in evolution and corresponds to R252 in the *Drosophila* WWE domain (Fig 2B). To examine the level of endogenous ADPR-Axin in *Iduna* mutants, ADPR-Axin was pulled down with wild type-WWE or R252A-WWE-mutant recombinant proteins (Fig 2C). This analysis revealed that *Iduna* mutants had a more than two-fold increase in ADPR-Axin in their midgut compared to wild type (Fig 2D-E). These suggest that Iduna promotes Axin degradation *in vivo*.

**Figure 2:**
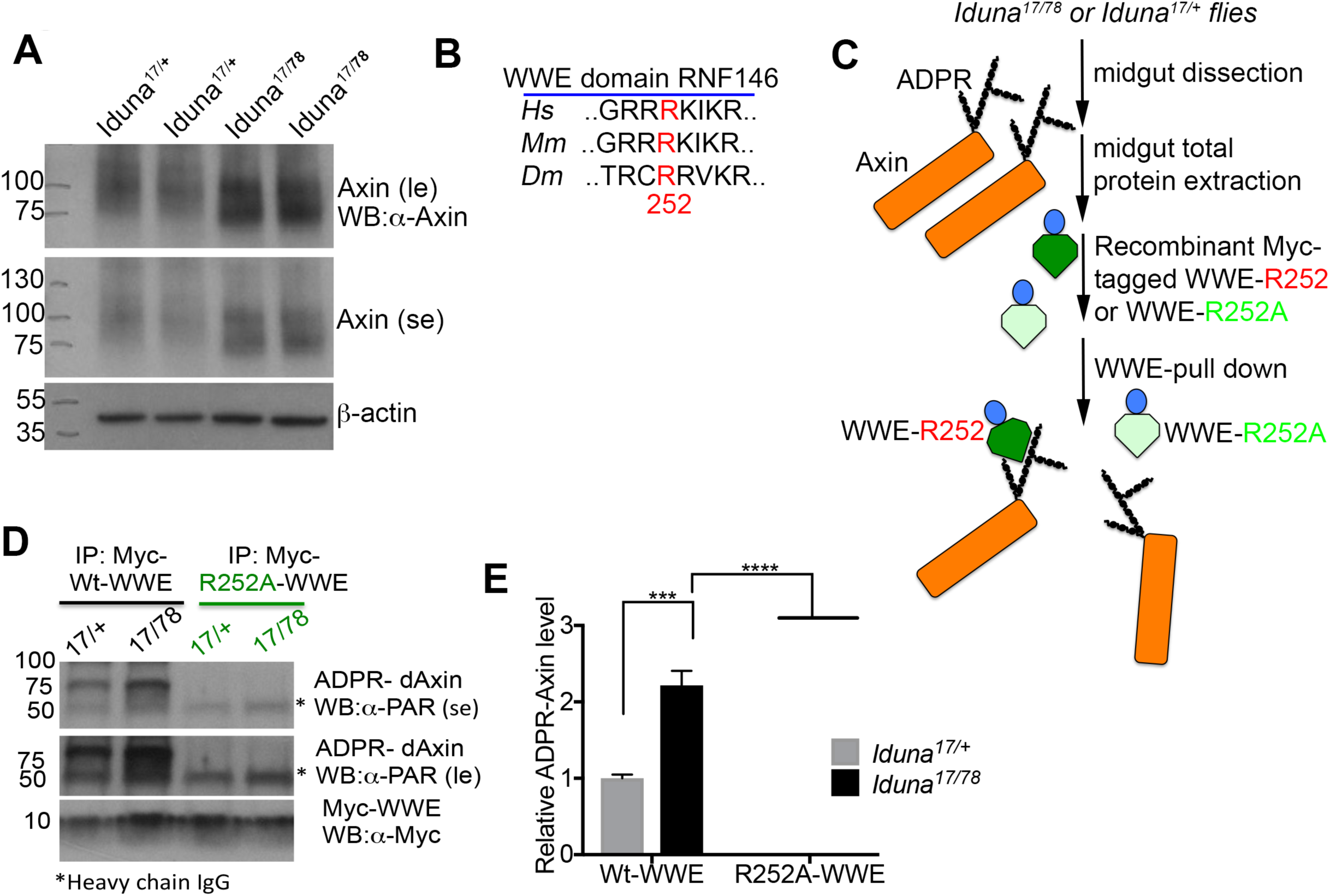
*Iduna* inactivation causes increased Axin protein levels in the midgut. **A-** *Iduna* mutant midguts have elevated Axin protein. Midguts of 7 day-old adult females expressing GFP-Axin under the temperature sensitive *Myo1A-Gal4* driver were dissected, lysed and analyzed by GFP immunoblotting. *Iduna* mutants had more Axin protein compared to the wild type. 20µg total intestinal lysates were analyzed by GFP immunoblotting and α-tubulin was used as a loading control. **B-** Loss-of-*Iduna* resulted in 2.2-fold GFP-Axin accumulation in the midgut. Western blot quantification was performed based on two independent experimental replicates, and protein levels were normalized to α-tubulin. **C-** Iduna recognizes ADP-ribosylated (ADPR) Axin via the R252 residue of its WWE domain. This residue corresponds to R163 in the mammalian Iduna ortholog and is conserved during evolution. Recombinant wild type and R252A mutants were used as biochemical sensors to pull down the ADPR-Axin from *Drosophila* midguts. Myc-tagged WWE proteins were expressed and purified from *Drosophila* S2R+ cells by immunoprecipitation. **D-** Inactivation of *Iduna* leads to accumulation of ADPR-Axin. Wild type myc-tagged-WWE protein pulled down ADPR-Axin. In contrast, the R252A mutant did not interact with modified Axin. Following IP, eluted proteins were analyzed with an α-PAR antibody. The 50kDa heavy chain IgG is indicated on the blot. **E-** *Iduna* inactivation results in 2.3-fold more ADPR-Axin protein in the midgut. Western blot quantification of two independent experimental replicates; ADPR-Axin levels were normalized to the control lines. Flies were fed with regular diet at 24-25°C. *p<0.001* is indicated as *** and *p<0.0001* was marked as ****.

To further understand the contribution of *Iduna* inactivation for both TNKS and Axin proteolysis in *Drosophila*, UAS-Flag-TNKS and UAS-GFP-Axin transgenes were mis-expressed under an eye-specific driver, *GMR*, in an *Iduna* mutant background (Fig S2A). To detect mis-expressed GFP-Axin and flag-Tankyrase levels, total proteins were extracted from five day-old male heads and analyzed by immunoblotting (Fig S2C and E). We found that *Iduna* mutants had 2.5-fold more mis-expressed GFP-Axin protein when compared to the control (Fig S2D). These mutants had 3.5-fold more ectopic expressed flag-tagged Tankyrase as well (Fig S2F). When we examined the eye morphology, GFP-Axin mis-expression did not cause obvious eye phenotype (Fig S2A).

On the other hand, mis-expressed flag-tagged Tankyrase led to rough eyes. This phenotype was more severe when *Tnks* was mis-expressed in *Iduna*^*-/-*^ homozygous mutants compared to *Iduna*^*-/+*^ heterozygous animals. (Fig S2B). Recently, it was also reported that mis-expressed Tankyrase promotes apoptosis in the *Drosophila* eye due to the activation of JNK signaling (Feng et al., 2018).

In order to examine whether Axin is a target for Iduna-mediated degradation, we also mis-expressed a UAS-GFP-Axin transgene under the enterocyte specific temperature sensitive *Myo1A-Gal4* driver (Fig S3A) and saw 2-2.5 fold more Axin in *Iduna* mutants compared to controls (Fig S4B). To investigate the cellular levels of Myo1A driven GFP-Axin in the enterocyes, we examined *FRT80B, Iduna* mutant clones and found that mutant enterocyte clones had more GFP-Axin when compared to their neighboring cells (Fig S4C). Taken together, these observations suggest that Iduna plays a role in promoting the degradation of both Axin and TNKS.

### Iduna is required to control the proliferation of intestinal progenitors in the *Drosophila* midgut

Attenuations in the Wingless pathway cause over-proliferation of stem cells in the *Drosophila* midgut. For instance, inactivation of *Tcf, arr, armadillo, dsh,* and *pygo* leads to suppression of Wingless signaling, which in turn causes more stem cell division (Kramp et al., 2002; Wang et al., 2016a and 2016b; Tian et al., 2016). On the other hand, *Apc* and *Tnks* mutations cause elevation of Axin, reduce wingless signaling and mitosis of stem cells in *Drosophila* (Wang et al., 2016a and 2016b; Tian et al., 2016). Hence, the Wingless signaling pathway is required to control intestinal stem cell proliferation in *Drosophila* (Xu et al, 2011; Cordero et al., 2012; Tian et al., 2016). Since *Iduna* mutants have elevated Axin level, we considered that *Iduna* inactivation may cause aberrant proliferation of stem cells in the *Drosophila* midgut. Just as the mammalian intestine (Korinek et al., 1998), the *Drosophila* midgut has intestinal stem cells (ISCs) which give rise to all intestinal compartments (Micchelli et al., 2005; Ohlstein et al., 2005). ISCs give rise to two types of daughter progenitor cells: undifferentiated enteroblasts (EBs) and pre-enteroendocrine cells (Pre-EE). EBs and pre-EEs differentiate into enterocytes (ECs) and enteroendocrine (EEs) cells, respectively (Ohlstein et al., 2005; Xu et al., 2011) (Fig S4A). Stem cells can be distinguished from enterocytes by their cell size and marker proteins (Ohlstein and Spadling 2006; Xu et al., 2011). Stem cells are small, express cell membrane-associated Armadillo, and lack of nuclear Prospero (Fig S4B). In contrast, nuclear Prospero staining is a marker of small-sized differentiated enteroendocrines (Fig S4B).

ISCs are also marked by the expression of *escargot (esg),* a transcription factor whose GFP reporter allows tracing of stem and progenitor cells during development (Ohlstein et al., 2005) (Fig S4B). Using the *esg>GFP* marker, we first analyzed 9 day-old female flies which were fed with a 5% sucrose diet for seven days at 28°C and saw an approximately twofold increase in the numbers of escargot positive ISCs/progenitors in the midgut of *Iduna* mutants compared to controls (Fig 3A-B). *Iduna* inactivation increased the numbers of Arm^+^/Pros^-^ stem cells in the midguts (Fig 3C) upon nutrient deprivation.

**Figure 3:**
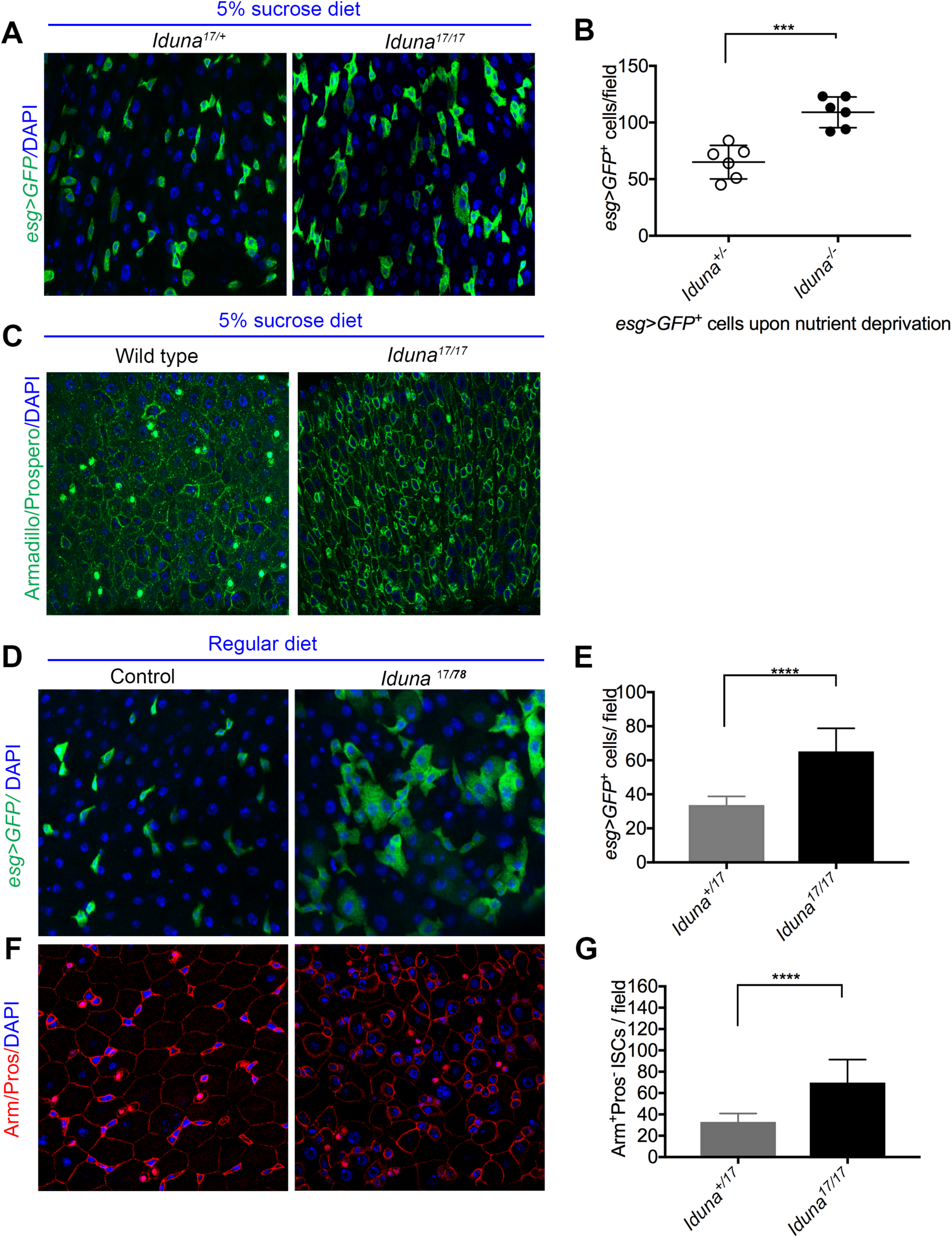
*Iduna* mutants have increased numbers of intestinal stem and progenitor cells in their midgut. **A-B-** Inactivation of *Iduna* promotes the proliferation of the esg-GFP labeled progenitor cells in the *Drosophila* midgut. Intestinal stem and progenitor cells were visualized with a *esg>GFP* reporter construct. **C-** Quantification of *esg>GFP*^*+*^ stem cells and progenitors from adult flies of indicated genotypes. **D-** In wild type, 25-30% of posterior midgut cells are stem cells, as assessed by *esg>GFP* expression. On the other hand, 55-60% of the total cell population in *Iduna* mutants expressed the stem marker *esg>GFP*, representing a greater than two-fold increase. **E-F-** *Iduna* mutants have more Arm^+^/Pros^-^ intestinal stem cells in the midgut. ISCs and enterocytes were distinguished by their cell size, high level of membrane-associated Armadillo, and lack of nuclear Prospero staining. In contrast, small-sized differentiated enteroendocrines were recognized by nuclear Prospero staining. Posterior midguts were analyzed by confocal microscopy following staining for anti-Armadillo, Prospero and DAPI. **G-H** Quantification of Arm^+^/Pros^-^ ISCs from adult flies of indicated genotypes. The midguts of seven day-old adult females were dissected and analyzed by confocal microscopy. For the consistency, posterior midgut R5 region was analyzed in this study. *Iduna*^*17/+*^ flies were used as control. Flies were fed with regular diet at 24-25°C. n>12 from each genotype. *p<0.001* was indicated as *** and *p<0.0001* was marked as ****.

To test whether the increased number of ISCs was dependent on nutrient deprivation, we examined midguts of seven-day old female mutants and controls on regular diet. We saw again an approximately twofold increase in the numbers of both *escargot>*GFP positive (Fig 3D-E) and Arm^+^/Pros^-^ stained (Fig 3F-G) stem cells-progenitors under these conditions. Therefore, increased ISC numbers in *Iduna* mutants are independent of diet.

To exclude the possibility that *Iduna* mutant flies raised on regular diet had reduced nutrient uptake, we monitored fly feeding by an acid blue 9 colorimetric assay (Mattila et al., 2018). We noticed no decrease in food intake in *Iduna* mutants kept on regular diet at 24-25°C compared to controls (Fig S1E). These results show that *Iduna* inactivation promotes the numbers of midgut stem cells independent of diet and food intake. Finally, we analyzed the midgut cell composition in *Iduna* mutant and control flies. We observed a slight increase in the total midgut cell number of *Iduna* mutants (Fig S4C). However, there were no significant differences in the number of EC and EE cells (Fig S4D-E). Collectively, these observations indicate that *Iduna* inactivation selectively affects ISC numbers.

The observed increase in stem cell numbers could be the result of aberrant stem cell proliferation or inhibition of their differentiation. To distinguish between these possibilities, we first assessed cell proliferation by dissecting 7 day-old mutant or wild type females. Following an hour EdU-labeling of the dissected midguts, we observed that *Iduna* mutants had more EdU-positive cells (Fig 4A-C). Moreover, phospho-Ser-Histone H3 (pH3) immunostaining (Fig 4D-E) also revealed a significant increase in pH3^+^ mitotic cells in the midgut of seven-day old female *Iduna* mutants (Fig 4D-F). These findings suggest that stem cells undergo increased proliferation in the midgut of *Iduna* mutants. To address whether there was an inhibition of differentiation in *Iduna* mutants, we generated *FRT80B, Iduna* mutant clones (Theodosiou et al., 1998). We found that ECs and EEs were present in the five-day old female mutant clones, demonstrating that Iduna was not essential for differentiation of ISCs into daughter cells (Fig S4F-G).

**Figure 4:**
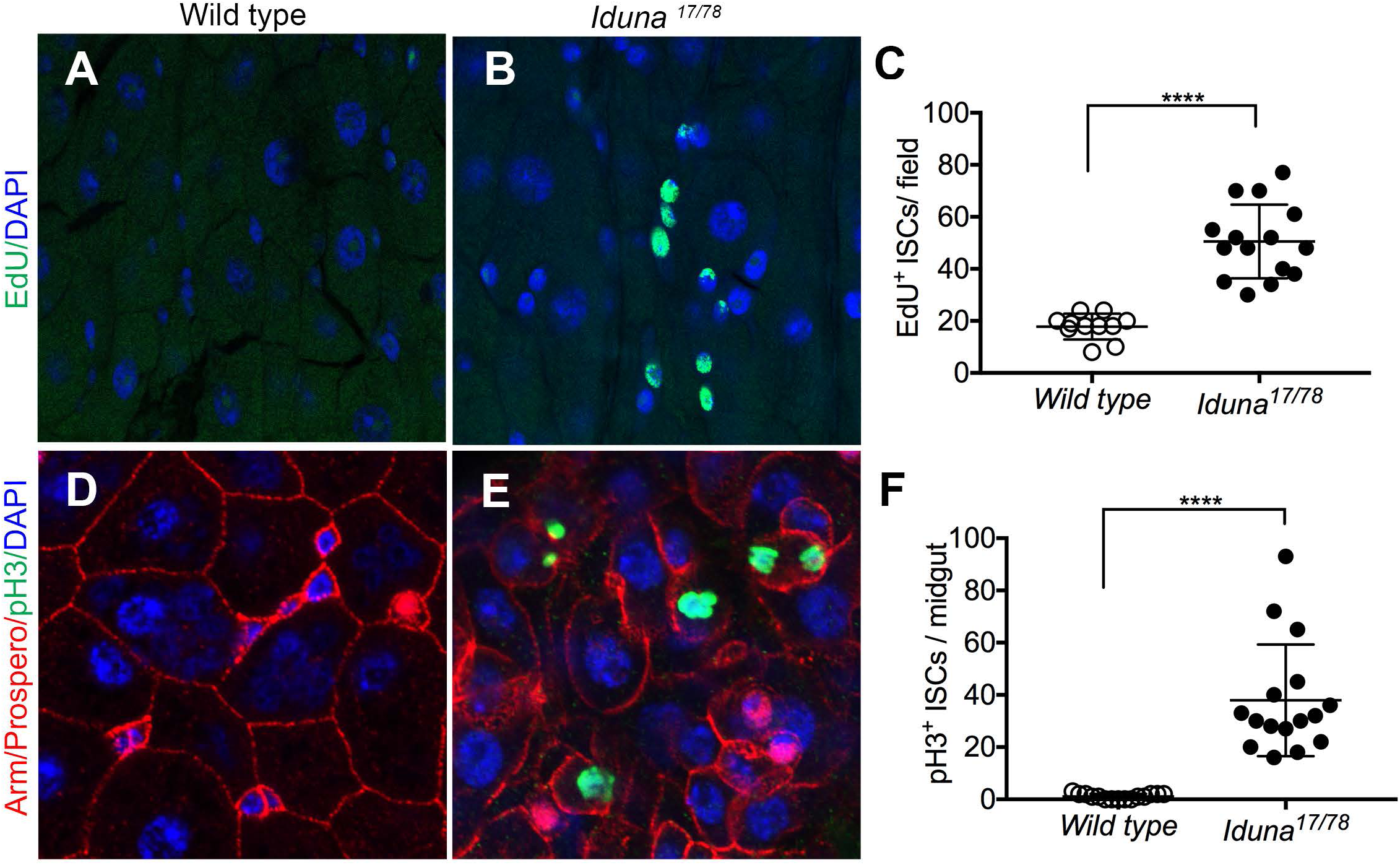
*Iduna* inactivation enhances proliferation of intestinal stem cells. **A-B-** Genetic depletion of *Iduna* leads to over-proliferation of intestinal stem cells in the midgut. EdU was used as a proliferation marker and administrated for 1h at 25°C. 7 day-old mutants or wild type female flies were dissected for analysis. Following fixation, EdU incorporation was analyzed by confocal microscopy. **C-** Increased numbers of EdU^+^ stem cells were seen in *Iduna* mutants, indicating increased cell proliferation. Posterior midguts were analyzed for quantification. **D-E-** *Iduna* mutants display elevated phospho-Ser-Histone 3 (pH3)-positive ISCs**. F-** Iduna inactivation leads to an increase of pH3^+^ mitotic stem cells in the midgut. Quantification of pH3^+^ proliferating cells was done in the whole midgut. 7 day-old mutants or wild type female flies were examined. Flies were fed with regular diet at 24-25°C. n>12 from each genotype. *p<0.0001* was marked as ****.

### Regulation of Axin proteolysis by Iduna is necessary for normal ISC proliferation

One possible mechanism by which Iduna may control the proliferation of ISCs in the *Drosophila* midgut is through modulating the concentration of Axin. To determine whether a reduction of the elevated Axin levels could reduce ISC numbers in *Iduna* mutants, they were recombined with *Axin* mutants and then crossed again with *Iduna* mutants to generate flies that were homozygous mutant for *Iduna*^*-/-*^ and heterozygous for *Axin*^+/-^. Strikingly, a reduction of the *Axin* gene dosage by 50% restored ISC numbers to wild type levels in *Iduna* mutants (Fig 5A). Compared to seven day-old female controls, *Iduna* mutants had an approximately twofold increase in the number of Arm^+^/Pros^-^ as well as pH3^+^ mitotic stem cells (Fig 5B-C). On the other hand, reducing the *Axin* gene dosage by 50% in an *Iduna*-Null background yielded numbers of ISCs and the pH3^+^ stem cells comparable to seven day-old wild type females. These results suggest that small changes in the levels of Axin have profound effects on stem cell number, and that regulation of Axin degradation by Iduna is necessary for normal ISC proliferation.

**Figure 5:**
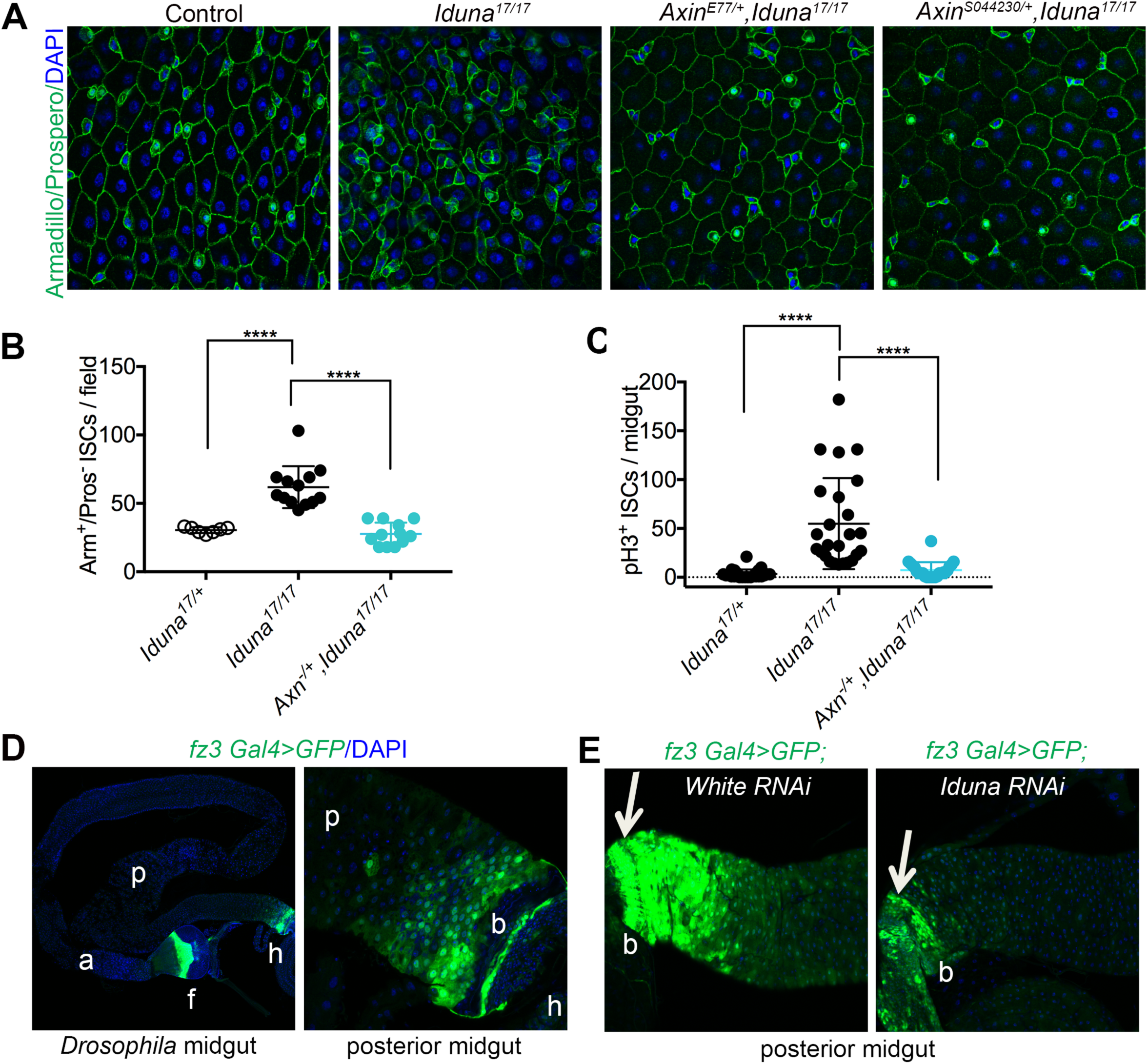
A 50% reduction of *Axin* restores ISC numbers in the *Drosophila* midgut. **A-** Reducing the *Axin* gene dosage by half restores the number of Arm^+^/Pros^-^ ISCs. *Axin* mutants *AxinS044230* and *AxnE77* were recombined with the *Iduna*^*17*^ mutant allele. *AxinS044230* is a complete *Axin*-Null mutant, and *AxinE77* is a loss-of-function truncation allele (Q406X). Midguts of 7d-old adult females of the indicated genotypes were dissected and analyzed by confocal microscopy following Armadillo, Prospero and DAPI staining. *Axin*^*+/-*^, *Iduna*^*17/+*^ served as control. Reducing the Axin gene dosage by 50% in *Iduna*^*17/17*^ mutants decreased the number of ISCs to normal levels. **B-** Quantification of ISC numbers. Reducing the *Axin* gene dosage by half fully suppressed the increased numbers of Arm^+^/Pros^-^ ISCs in *Iduna*^*17/17*^ Null-mutants. **C-** Reducing *Axin* gene expression suppressed the proliferation of ISCs in the *Iduna*^*17/17*^ Null-mutant. *p<0.0001* was marked as ****. **D-** *Frizzled 3 (fz3)* is a *wingless* target gene and a GFP-reporter construct was used here to visualize Wg-activity in the midgut (Buchon et al., 2013; Tian et al., 2016; Wang et al., 2016a, b). In wild type, *fz3>GFP* is highly expressed in a graded fashion in the foregut (f), the posterior midgut (p) as well as the posterior midgut-hindgut border, but not in the anterior midgut (a) or the hindgut proper (h). Higher magnification image of the *fz3>GFP* near the midgut-hindgut boundary. 7-day old female midguts were analyzed. **E-** *fz3-Gal4* driven *Iduna* depletion inhibits *wingless* activity. RNAi-mediated down-regulation of *Iduna* led to significant reduction of *fz3>GFP*; *white RNAi* served as a control. The arrow indicates the border between the posterior midgut and the hindgut (b). Three day-old female midguts were analyzed. Flies were fed with regular diet at 24-25°C.

We observed that *Iduna* mutants had 2-fold more Axin in the *Drosophila* midgut. This indicates that defects in Axin degradation may cause over-proliferation of stem cells due to inhibition of wingless signaling. Therefore, we analyzed a reporter for the Wingless pathway target gene, *frizzled-3* (*fz3*). It was previously reported that *fz3-RFP* reporter activity is high at the major boundaries between compartments (Buchon et al., 2013; Tian et al., 2016; Wang et al., 2016a, b). *Fz3-RFP* was strongly expressed in enterocytes at three distinct sites of the midgut: around R1a, R2c, and R5 (Buchon et al., 2013). Therefore, enterocytes are the primary sites of the Wingless pathway activation during intestinal homeostasis (Tian et al., 2016).

We analyzed 3 day-old *fz3-Gal4>GFP* expressing females and consistently observed that *fz3>GFP* was expressed in gradients in the foregut, posterior midgut, as well as the border between the posterior midgut and hindgut (Fig 5D). Here, we focused on the posterior midgut-hindgut border to investigate the effect of Iduna on wingless signaling. Upon *fz3-Gal4* driven RNAi-mediated *Iduna* depletion, we found that *fz3>GFP* activity decreased significantly (Fig 5E). We conclude that Iduna stimulates *wingless* activity in the posterior midgut by promoting degradation of Axin.

The proliferation of stem cells in the *Drosophila* midgut is regulated by intrinsic signals and also interactions with neighboring cells (Zhou et al., 2013; Tian et al., 2016). To further investigate whether the observed effects could reflect a cell-autonomous requirement of Iduna in stem cells, or alternatively a requirement in other cells of the midgut, *Iduna* was specifically targeted in enterocytes as well as midgut stem and progenitors cells by using the *Myo1A-Gal4* and *esg-Gal4* drivers, respectively (Fig 6A-B). We examined 7 day-old females expressing *Iduna* RNAi under the *Myo1A* or *esg* drivers. RNAi-mediated knock-down of *Iduna* in enterocytes caused a significant increase in Arm^+^/Pros^-^ stem cell numbers (Fig 6B). However, stem cell/progenitor cell-specific knock-down of *Iduna* did not affect either the stem cell numbers or mitosis in the midgut (Fig 6B-C). This suggests that *Iduna* inactivation causes stem cell over-proliferation by a non cell-autonomous mechanism, and that perhaps enterocytes are responsible for stem cell over-proliferation in *Iduna* mutants. To further test this idea, we ectopically expressed *Iduna* in enterocytes and investigated if this could suppress stem cell proliferation in *Iduna* mutants (Fig 6D). Indeed, consistent with this model, we saw that *Myo1A-Gal4* driven UAS-Iduna was able to restore normal numbers of stem cells and progenitors (Fig 6E-F). Taken together, our results indicate that Iduna plays a physiological role to regulate *wingless* signaling in enterocytes, which is critical for proper ISC proliferation.

**Figure 6:**
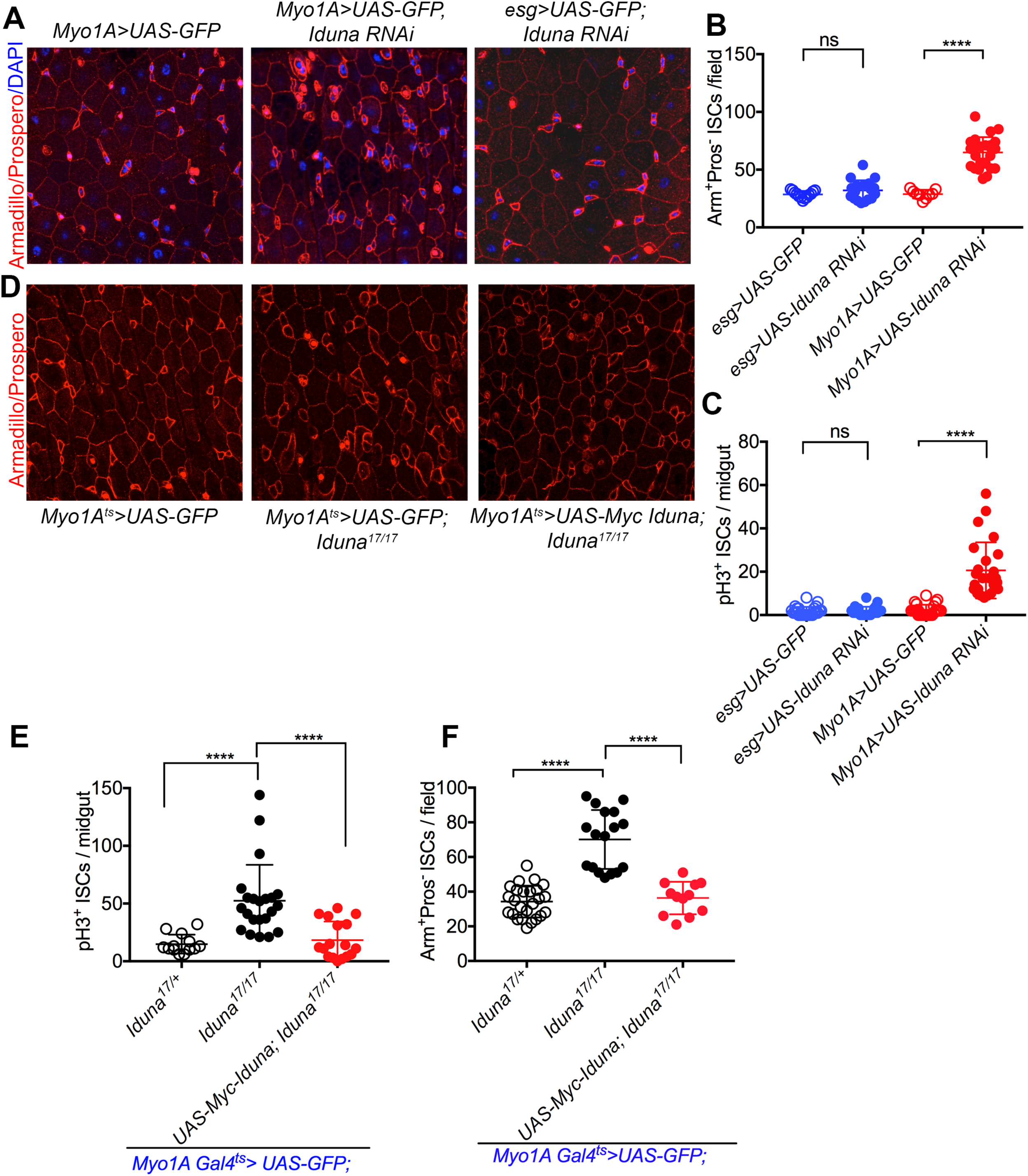
*Iduna* depletion in ECs leads to over-proliferation of ISC. **A-** Over-proliferation of ISCs in *Iduna* mutants is non-cell autonomous. RNAi-mediated *Iduna* knock-down was carried out in the enterocytes, stem cells and enteroblast cells using *Myo1A-Gal4* and *esg-Gal4* drivers, respectively. **B-** Knock-down of *Iduna* in enterocytes using the *Myo1A-Gal4* driver led to over-proliferation of Arm^+^/Pros^-^ ISCs. In contrast, no changes in ISC proliferation were observed upon down-regulation of *Iduna* in ISCs using *esg-Gal4*-driven *Iduna* RNAi. *Myo1A-Gal4>*GFP served as a control. **C-** EC-specific knock-down of *Iduna* increased the number of pH3^+^ progenitors. **D-** Ectopic expression of *Iduna* in ECs inhibits over-proliferation of ISCs. A UAS-Myc-tagged Iduna C/G transgene was generated to perform rescue experiments. **E-F-** Expression of the UAS-Myc-Iduna C/G transgene with the *Myo1A-Gal4* driver resulted in a reduction of the numbers of Arm^+^/Pros^-^ ISCs and pH3^+^ mitotic stem cells in the midgut of *Iduna* mutants. Flies were fed with regular diet at 24-25°C. 7 day-old female midguts were analyzed for ISC and mitotic markers. *p<0.0001* was marked as ****.

We found that *Iduna* mutants have increased mortality upon nutrient deprivation (Fig. 1C). Following 7 days on a 5% sucrose diet at 28°C, *Iduna* mutants had more *esg>GFP* positive cells in the midgut (Fig 3A-C). Therefore, we considered that under reduced nutrient diet, hyper-proliferation of midgut stem cells may be responsible for elevated mortality. To test this idea we first inactivated *Iduna* in enterocytes by expression of three different RNAi lines under the Myo1A driver. We found that RNAi-mediated *Iduna* depletion did not increase lethality compared with *white* RNAi (Fig S5A). There was also no significant change on the mean lifespan between *white* and *Iduna* RNAi expressing flies (Fig S5B). We also tested enteroblast specific *Iduna* depletion and again found no significant effects on longevity upon nutrient deprivation (Fig S5C). Finally, we expressed UAS-*Iduna* transgene *under Myo1A driver* in enterocytes to rescue the elevated mortality in *the* mutants. Whereas the *Iduna* transgene rescued the hyper-proliferation phenotype (Fig S5E), it failed to rescue the mortality of mutants on 5% sucrose diet (Fig S5D). These findings suggest that Iduna mortality is not caused by dysregulation of midgut stem cell proliferation and point to another role of *Iduna* in promoting survival under stress conditions.

### Depletion of *Iduna* promotes stem cell proliferation through the JAK-STAT pathway

In order to further investigate the mechanism by which Iduna affected ISC proliferation, we explored the function of additional signaling pathways implicated in this system. Because the JAK-STAT pathway has a well-known role in stem cell proliferation (Zeidler et al., 2000; Zoranovic et al., 2013; Zhou et al., 2013; Markstein et al., 2013), we looked for possible effects of *Iduna* mutants here. We analyzed the JAK-STAT pathway via 10x Stat-GFP reporter line in the midgut (Bach et al., 2007).

Under regular physiological conditions, Stat-GFP reporter expression was mainly seen in small sized cell populations in the midgut that appear to represent ISCs for several reasons (Fig S6). First, Prospero-positive enteroendocrine cells were negative for Stat-GFP (Fig S6A). Second, enterocytes stained with Armadillo were also not expressing the Stat-GFP reporter (Fig S6D). Finally, Delta-lacZ positive but Prospero-negative cells for the most part expressed Stat-GFP. However, a minor population of small sized cells was GFP positive but Delta-lacZ negative (white arrows, Fig S6B). These appear to be undifferentiated progenitors, such as enteroblasts. Seven day-old *Iduna* mutants had more Stat-GFP positive cells when compared to controls (Fig 7A, S7A-F). We also generated *FRT80B, Iduna* midgut mutant clones and observed that these clones had elevated JAK-STAT signaling (Fig S7A-B). To confirm elevated JAK-STAT signaling in *Iduna* mutant stem cells, we stained midguts from seven day-old females for Delta, a previously identified JAK-STAT pathway target gene (Jiang et al., 2009). We found that there was indeed more Delta protein in *Iduna* mutants, consistent with elevated JAK-STAT activity (Fig S7G).

**Figure 7:**
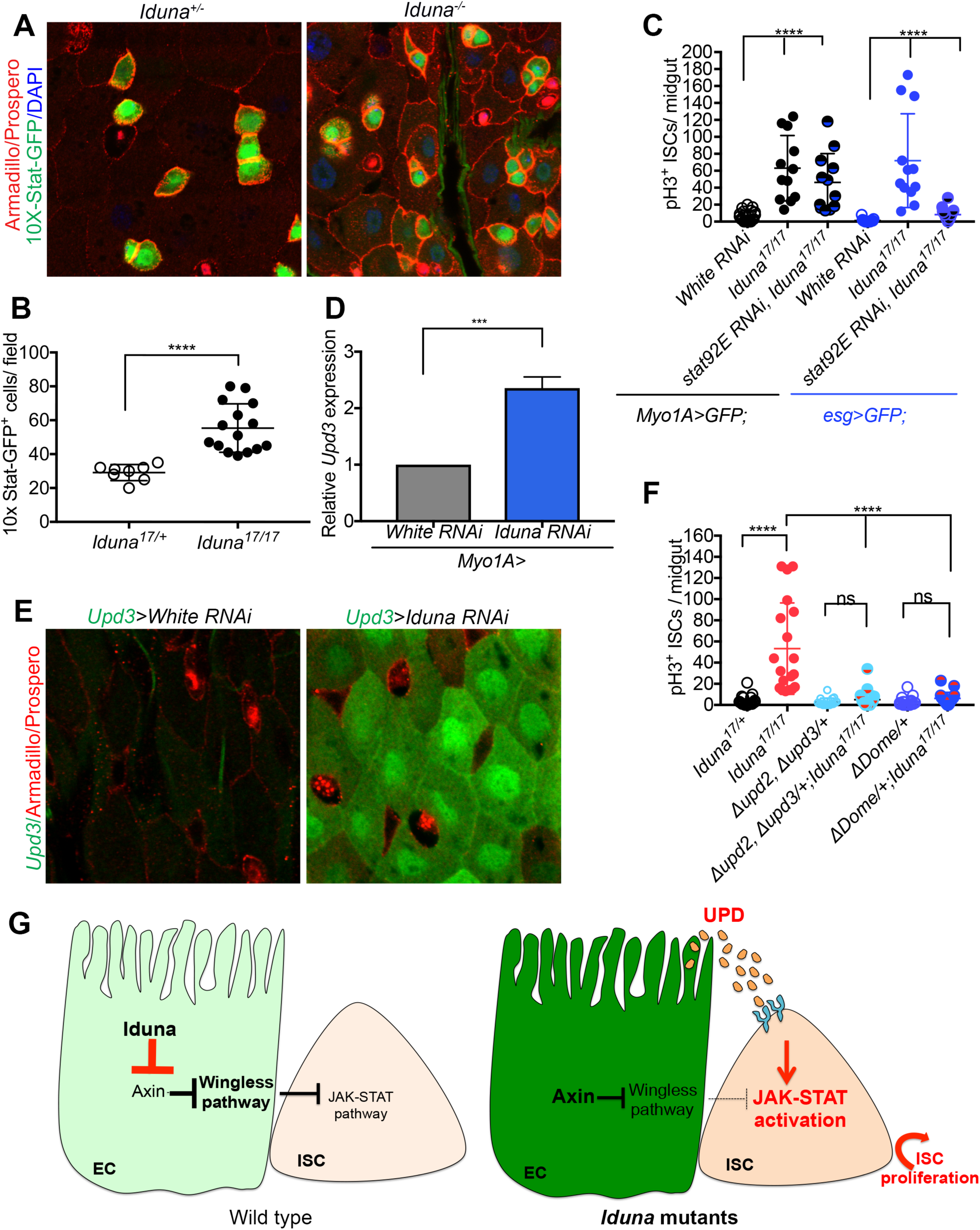
Loss of *Iduna* activates the JAK-STAT pathway non-cell autonomously to promote ISC proliferation. **A**-*Iduna* mutants have elevated Stat-GFP signaling in ISCs. *10X-Stat-GFP* is a reporter for STAT signaling activity. *Iduna* mutants displayed strongly increased GFP reporter. 7 day-old females were dissected and the posterior midguts were analyzed. **B-** Quantification of Stat-GFP expressing midgut cells. **C-** Knock-down of the *stat93E* transcription factor in ISCs and EBs blocks ISC over-proliferation in *Iduna* mutants. In contrast, RNAi-mediated depletion of *stat92E* in ECs did not affect proliferation of ISCs. 7 day-old female midguts were dissected and analyzed. **D-** *Iduna* depletion results in upregulation of *Upd3* mRNA expression in enterocytes. Myo1A driven *Iduna-RNAi* and *white RNAi* expressing 7 day-old females were dissected for their midguts. Total RNA was isolated and cDNA libraries were prepared. *Upd3* transcripts were amplified and analyzed by q-PCR. **E***-* RNAi-mediated *Iduna* down-regulation induces *Upd3>GFP* reporter activity. *Iduna* was knocked-down using RNAi driven by *Upd3-Gal4*, and *GFP* was used as a reporter for *Upd3* gene expression. Enterocytes were stained with α-Armadillo antibody. EEs and ISCs were negative for Upd3>GFP expression. *white RNAi* served as a control. 3 day-old female flies were dissected and their posterior midguts were analyzed by confocal microscopy. **E-** Reduction of either *Upd2 and Upd3* or their receptor *Domeless* suppresses over-proliferation of ISCs in *Iduna* mutants. Upon reduction of *Upd2* and *Upd3* gene dosage in *Iduna* mutants, we observed significantly less mitotic stem cells in *Iduna* mutants, comparable to WT levels. Likewise, a 50% reduction of *Domeless* resulted in suppression of ISC over-proliferation in *Iduna* mutants. On the other hand, these reductions in gene dosage of *Upd2, Upd3* and *Dome* did not affect mitosis of ISCs in a WT background. 7 day-old female midguts were quantified by pH3^+^ staining. Flies were fed with regular diet at 24-25°C. n>12 from each genotype. *p<0.0001* was marked as ****. G-Model for the role of *Iduna* in the regulation of intestinal stem cell proliferation. Our model suggests that inactivation of *Iduna* causes Axin elevation which in turn decreases wingless signaling activation in enterocytes, and increases secretion of UPD cytokines from these cells. These cytokines activate JAK-STAT signaling through the Dome receptor on neighboring ISCs and thereby induce ISC proliferation in the *Drosophila* midgut.

To test whether activation of JAK-STAT signaling was responsible for aberrant ISC proliferation, we knocked down *stat92E*, a transcription factor in the JAK-STAT pathway, in enterocytes as well as in stem cells and progenitors. We did not detect dramatic changes in the numbers of mitotic cells when *stat92* transcription factor was depleted in enterocytes (Fig 7B). Interestingly, knock-down of *stat92E* in midgut stem and progenitors cells was sufficient to suppress their increased cell division (Fig 7B). Collectively, these observations suggest that *Iduna* inactivation causes decrease in *wingless* signaling in enterocytes which in turn causes elevated JAK-STAT signaling in midgut stem cells and thereby results in their over-proliferation.

Our observations raise the question of how enterocytes signal ISC proliferation. One possibility is that enterocytes secrete a factor activating the JAK-STAT pathway in stem cells. The JAK-STAT pathway can be activated by cytokines, such as *unpaired* (UPD, UPD2, UPD3) in the *Drosophila* midgut (Ghiglione et al., 2002; Zhou et al., 2013). Upd3 is produced in differentiated ECs and in differentiating enteroblasts (Zhou et al., 2013). Therefore, we explored the possibility that *unpaired* cytokines could mediate stem cell over-proliferation in *Iduna* mutants. For this purpose, we first inactivated *Iduna* with the *Upd3-Gal4* driver and found that RNAi-mediated knock-down of *Iduna* resulted in a significant increase of *Upd3>GFP* reporter expression in the midgut (Fig 7C-D, Fig S8A). *Upd3>GFP*-positive cells were mainly enterocytes, and not EEs or ISCs (Fig 7E, Fig S8B-C). We then knocked down *Iduna* in enterocytes and performed a q-PCR to test whether *Iduna* depletion in enterocytes induced unpaired expression. We detected that enterocyte specific *Iduna* inactivation resulted in elevated *Upd3* gene expression compared to *white* RNAi (Fig 7D). To suppress the over-proliferation of midgut stem cell in *Iduna* mutants, we finally reduced *Upd2* and *Upd3* gene dosages. Strikingly, we found that heterozygosity in Δ*Upd2-Upd3* fully suppressed ISC proliferation in *Iduna* mutants (Fig. 7E). Secreted Unpaired proteins bind to the Domeless receptor on ISCs (Ghiglione et al., 2002). Therefore, we tested whether decreasing Domeless levels could also suppress stem cell over-proliferation in *Iduna* mutants. Again, this prediction was experimentally confirmed (Fig 7E). We conclude that inactivation of *Iduna* causes decrease in wingless signaling in enterocytes, which in turn leads to increased secretion of UPD2-3 from these cells to stimulate over-proliferation of ISCs through the JAK/STAT pathway (Fig 7F).

## Discussion

In this study, we investigated the *in vivo* function of *Iduna* and identified a critical role of this enzyme for the control of stem cell proliferation in the *Drosophila* midgut. It was previously shown that mammalian Iduna is an unusual E3-ubiquitin ligase that specifically binds to and poly-ubiquitylates ADP-ribosylated substrates to promote their rapid degradation by the proteasome. However, the physiological function of Iduna remains largely unclear. Here, we generated *Drosophila* Null-mutants and used them to show that Iduna has an important *in vivo* function for the degradation of ADP-ribosylated TNKS and Axin to control stem cell proliferation. In particular, we focused on the role of *Iduna* in the *Drosophila* midgut. We found that *Iduna* inactivation caused a slight but significant increase in Axin protein levels in enterocytes, which in turn caused over-proliferation of intestinal stem cells. This non-cell autonomous effect on stem cell proliferation was depended on UPD2-UPD3 cytokines that are secreted from enterocytes. These findings suggest a model in which loss of Iduna function, which decreases the Wingless pathway activity due to elevated Axin levels in enterocytes, which in turn causes increased secretion of UPD2-3 resulted from these cells to activate the JAK-STAT pathway in ISCs. Importantly, a 50% reduction in *Axin* gene dosage blocked the over-proliferation of stem cells in *Iduna* mutants, demonstrating the necessity to tightly regulated Axin levels in this system. Whereas many other cell types appear to tolerate fluctuations in the amount of Axin protein, proper wingless signaling in the *Drosophila* midgut appears to critically depend on restricting Axin levels by Iduna. The activity of Iduna depends on binding to ADP-ribosylated substrates via its WWE domain. Recognition and binding to its ADP-ribosylated target proteins change the structural confirmation of Iduna. Subsequently, Iduna is activated to ubiquitylate its targets for proteasome-mediated degradation. It was previously reported that TNKS forms a tight complex with Iduna to control the proteolysis of target proteins (DaRosa et al., 2015). We could not detect any obvious morphological differences between *Iduna* mutants and wild type. Although this may seem somewhat surprising, it is consistent with inactivation of *Tnks* in *Drosophila,* which also causes no overt abnormalities (Feng et al., 2014; Wang et al., 2016a and 2016b; Yang et al., 2016). Like for *Iduna, Tnks* mutants have no obvious effects on wing development and the expression of *wingless* target genes in larval wing discs, despite the fact that Axin levels are increased (Feng et al., 2014; Wang et al., 2016a and 2016b; Yang et al., 2016). Our interpretation of these findings is that most tissues can tolerate relative modest (2-3-fold) changes of Axin. For example, it appears that a greater than 3-fold increase of endogenous Axin is required for functional consequences of altered wingless signaling in *Drosophila* embryos (Yang et al., 2016) and 3-9-fold changes are needed in wing discs (Wang et al., 2016a). On the other hand, the *Drosophila* midgut appears much more sensitive to reduced Wingless signaling.

A recent study demonstrated that inactivation of *Drosophila Tankyrase (Tnks)* also led to increased Axin protein accumulation in the *Drosophila* midgut and promoted ISC proliferation as well (Wang et al., 2016). These results are consistent with previously reported cell-based studies suggesting that Iduna mediates Tankyrase-dependent degradation of Axin and thereby positively regulates Wnt signaling (Huang et al., 2009; Zhou et al., 2011; Croy et al., 2016; Callow et al., 2011). On the other hand, it is somewhat surprising that inactivation of two highly diverse types of enzymes, *Tankyrase*, a poly-ADP-ribose polymerase versus *Iduna*, a ubiquitin E3 ligase, produces remarkably similar phenotypes. Both Tnks and Iduna have many other targets outside the Wnt-pathway, and based on biochemical observations it has been proposed that they may play roles in DNA repair, telomere length, vesicle trafficking, Notch-signaling, centrosome maturation, neuronal protection and cell death (Bai et al., 2012; Gibson et al., 2012; Riffel et al., 2012). However, *Iduna* mutant flies are viable and do not show any obvious defects under normal growth conditions. This indicates that the major non-redundant physiological function of both Tnks and Iduna in *Drosophila* is to regulate Wingless-mediated intestinal stem cell proliferation, and that it provides physiological evidence for the idea that the function of both proteins is indeed tightly coupled. In addition, our study identifies a role of Upd/Dome in this pathway. These results may also have implications for the regulation of this highly conserved pathway in mammals. For example, conditional inactivation of *Iduna* in mouse bones leads to increased numbers of osteoclasts and inflammation (Matsumoto et al., 2017a). In this system, down-regulation of *Iduna* leads to accumulation of Axin1 and 3BP2. This, in turn, attenuates β-catenin degradation and activates SRC kinase, respectively, thereby promoting the release of inflammatory cytokines in the bone (Matsumoto et al., 2017a). On the other hand, *Iduna* depletion reduces proliferation of osteoblasts and promotes adipogenesis in the mouse skeleton (Matsumoto et al., 2017b). Despite the obvious differences between mammalian bone and the *Drosophila* midgut, both systems show overall striking similarities in the use of TNKS/Iduna to restrict Axin levels to achieve proper levels of Wnt/β-catenin signaling during tissue homeostasis. Finally, our study also indicates that Axin may have a more general function as a scaffold protein to recruit multiple proteins to permit a crosstalk with other pathways to modulate Wnt/β-catenin signaling.

## Material and Methods

### Fly stocks

Flies were kept at a 12-hour light/dark cycle. All crosses were performed at 22-25°C unless stated otherwise. The following fly stocks were used for this study (Bloomington Drosophila Stock Center (BDSC) and Vienna Drosophila Resource Center (VDRC) number (#) given in parentheses):

The stocks used in here: Df(3L)Exel6135 (BDSC# 7614), Df(3L)ED228 (BDSC# 8086), Df(3L)ED229 (BDSC# 8087), esg-Gal4, UAS-GFP (a gift of Dr. Norbert Perrimon, Micchelli et al., 2006), esgK606 (a gift of Dr. Norbert Perrimon, Micchelli et al., 2006), *10X STAT-GFP*, Bach et al., 2007), UAS-GFP-Axin (BDSC# 7224), FRT82B, *Axin044230* (a gift of Dr. Wei Du, Hamada et al., 1999), FRT82B, *AxinE77* (a gift of Dr. Jessica Treisman, Collins et al., 2000), Myo1A-Gal4, tub-Gal80ts, UAS-GFP (a gift of Dr. Norbert Perrimon, Micchelli et al., 2006), Upd3-Gal4, UAS-GFP (a gift of Dr. Norbert Perrimon, Markstein et al., 2013), ΔUpd2/3 (BDSC# 129), ΔDome (BDSC# 12030), UAS-*stat92E* RNAi (BDSC# 26889), UAS-CG8786/*dIduna* RNAi#1 (BDSC# 40882), UAS-CG8786/*dIduna* RNAi#2 (VDRC#43533), UAS-CG8786/*dIduna* RNAi#3 (VDRC#36028), UAS-CG8786/*dIduna* RNAi#4 (VDRC#36029), and *white* RNAi (BDSC#33623), fz3-Gal4 (BDSC#36520). The rest of *Drosophila* lines, which were studied here, were obtained from Steller Lab stocks. Oregon R flies were used as control and only adult female flies were analyzed in this study.

### *Drosophila* egg collection

A 10mm^2^ apple-agar plate was set up with embryo collection cage to provide a substrate for egg laying. Prior to adding the plate, a small quantity of yeast paste was smeared onto the center of the apple-agar. To provide moisture, water soaked tissue paper was layered under embryo collection cages. 10-15 day old adult flies were collected to the cage, which were then placed into a fly incubator for 4 hours. Then, laden eggs were counted and 50 of them were plated into one corner of 10cm^2^ apple-agar plates, in which had a straight yeast paste smear at the center. Agar plates finally were incubated in the incubator. After 24h, hatched eggs were counted.

To analyze larval development, hatched 1^st^ instar larvae were counted and placed into a fresh yeast paste containing agar plate until they were reached to 3^rd^ instar larvae. After counting, larvae were placed into regular food containing vials. They were counted two rounds when they pupated and enclosed.

### 5% sucrose diet

5mm^2^ Whatman filter papers were soaked with 1ml 5% sucrose solution and placed into the empty vials. 5% sucrose solution was used as reduced nutrient diet. Enclosed adult females were collected at 24-25°C. When they were 2 day-old, their regular diet was replaced to 5% sucrose diet at 28°C. 20 of two day-old wild type or *Iduna* mutant female flies were grouped and kept on 5% sucrose solution-soaked filter paper containing vials at 28°C. Following the fly count, death flies were removed and 1ml 5% sucrose-embedded filter papers were replaced with a new one everyday.

### Food intake measurement

Female flies of *Iduna* mutant and Oregon R were collected after they enclosed. Before measuring food intake, the flies were kept on the regular food for 6 days. The flies then transferred to the regular food supplemented with 0.5% (w/v) Acid Blue 9 (erioglaucine disodium salt, Sigma 861146) for 4 hours. Quandruplicates of 5 flies per sample were then homogenized in 250µl 1xPBS and cellular debris was removed by centrifugation. Food intake was quantified by measuring the absorbance of the supernatant at 630nm and normalized to the wet weight of the flies.

### *CG8786/dRNF146/dIduna* CRISPR/Cas9 editing

We used the CRISPR optimal target finder website (tools.flycrispr.molbio.wisc.edu/targetFinder) to identify an appropriate guide RNA (gRNA) target sequence within *dIduna* (Granz et al., 2013 and 2014). We purchased the forward 5‵-GTCGCTAGCTGCAATCTGCTCTG-3‵ and reverse 5‵-AAACCAGAGCAGATTGCAGCTAG-3‵ oligos (IDT, Inc.) annealed, and followed the protocol; from Port et al., 2014 to clone the annealed oligos into pCFD3-dU6:3-gRNA plasmid (Addgene, plasmid# 49410, Port et al., 2014). Transformants were verified via Sanger sequencing (Genewiz, Inc.). The gRNA plasmid was injected into 300 embryos of custom *vasa*-Cas9 *Drosophila* (BestGene, Inc.). The injection was yielded 89 G_o_ progeny, and we established 70 individual fly lines, a couple of which could possibly have the *Iduna* loss-of-function mutations.

### Isolation of the *Iduna* mutants and genetic mapping of *Iduna* – loss-of-function mutations

Total DNA was isolated from L3 larvae or 5-days old adults of *Iduna* homozygous mutants and control sequencing strain using the Roche genomic DNA extraction kit (Roche). To confirm the mutant line, PCR fragments were amplified with specific primers (forward primer 5‵-CAGCCCGAGCTGGTCATACTCAG-3‵, reverse primer 5‵-CGGCTTTCTGGGCTACCTAC-3‵) binding within 5‵ UTR of *Iduna* and within the coding region of the gene. To identify mutation site, the entire coding region was PCR amplified and PCR products were sent for DNA sequencing.

### Cloning and generation of UAS-CG8786 transgenic *Drosophila*

Adult flies were directly homogenized in 1ml TRIzol (Life Technologies) and total RNA was isolated according to the manufacturer’s protocol. cDNA library was prepared from 5 µg total RNA, by using oligo(dT) amplification and the Superscript III First Strand synthesis kit (Invitrogen). cDNA library was used to amplify the *Iduna* transcripts with the primers (forward 5‵-ATGTCGCAACAGCGCTCCACAG-3‵; *Iduna* B isoform reverse primer 5‵-TCAGTAGAGCTTTAGGTATACC-3‵; *Iduna* C/G isoform reverse primer 5‵-TCAGTAGAGCTTTAGGTATACCG-3‵). Amplified *dIduna* transcripts were cloned into pUAST (DGRC, Drosophila Genomic Resource Center) and pAc5.1 (Thermo Scientific) vectors by considering the appropriate restriction digestion sites. Following the bacterial transformation, all of the cloned genes were sequenced. To generate UAS-CG8786 transgenic *Drosophila,* myc-tagged pUAST-CG8786/dIduna plasmid was injected into w1118 embryos (Best Gene, Inc.) We obtained successful transgenic lines.

### Total RNA isolation, cDNA synthesis and Q-PCR

Posterior midguts of 7 day-old adult flies were directly homogenized in 1 ml TRIzol (Life Technologies) and total RNA was isolated according to the manufacturer’s protocol (miRNeasy mini kit, QIAGEN). cDNA library was prepared from 5 µg total RNA, by using oligo(dT) amplification and the Superscript III First Strand synthesis kit (Invitrogen). cDNA library was used to amplify *Upd3* and *Rp32l* transcripts with the forward 5‵-AGGCCATCAACCTGACCAAC-3‵, reverse 5‵-ACGCTTCTCCATCAGCTTGC-3‵ and forward 5‵-CCCAAGGGTATCGACAACAGA-3‵, reverse 5‵-CGATCTCGCCGCAGTAAAC-3‵ primers, respectively. Those primers were designed via the online tool of DRSC/TRiP Functional Genomics Resources, Harvard Medical School and purchased from IDT, Inc.

### Cloning and generation of the wild type and mutants UAS-Flag-Tnks transgenic *Drosophila*

We previously described *Drosophila* TNKS (Park and Steller, 2013) and its ORF was cloned into pUAST vector from pcDNA3.1-Flag-TNKS. To generate UAS-Flag-TNKS transgenic *Drosophila,* Flag-tagged pUAST-TNKS plasmid was injected into w1118 embryos (Best Gene, Inc.) We obtained successful transgenic *Drosophila* lines and those were utilized in conjunction with tissue-specific *Gal4* drivers.

### Clone analysis and RNAi experiments

Mutant clones were utilized to generate mitotic clones. 2^nd^ instar larvae were subjected to an hour heat shock in a 37°C water bath per day until they reached the pupa stage and maintained at 24°C. 3-days old adult females were analyzed.

For RNAi experiment, crosses were performed at 24°C and the progeny of the desired genotypes were collected on the day of eclosion and maintained at 24°C for 7 days before dissection. In the case of using temperature sensitive driver, eclosed virgin females were collected and kept at 29°C for 7 days for intestine dissection.

### Cell culture

S2R+ cells were maintained at 25°C in supplemented Grace’s Insect Medium (supplemented with 10% heat inactivated FBS, 100U/ml penicillin, 100µg/ml streptomycin) in spinner flasks.

### Development of polyclonal antibodies

Full-length GST-tagged-Iduna C/G protein was expressed and purified from BL21 DE3 *E.coli* strain. Polyclonal antisera were generated in two guinea pigs (Cocalico, Inc.). For Western blot analysis, serum was used in 1:1000 dilutions.

### Western blot analysis

50-100µg dissected tissues or total larvae/flies were lysed in lysis buffer [50mM HEPES-KOH pH 7.4, 150mM NaCl, 0.05% Triton-X100, complete EDTA-free protease inhibitor cocktail (Roche)] using a 1 ml tissue grinder. Lysates were cleared by centrifugation at 13,000 g for 20 min at 4°C. Protein concentrations of supernatants were determined by BCA assay (Pierce). 1µg/µl lysate was prepared with 3X sample buffer in 100µl total volume (200mM Tris-HCl, pH 6.8, 200mM dithiothreitol (DTT), 8% SDS, 24% glycerol, 0.04% bromophenol blue), heated at 95°C for 10 min and samples were separated by SDS-PAGE for 1 h at 120V, by using standard 1X SDS Tris base-glycine running buffer. Proteins on the gels were blotted onto a PVDF membrane, in a 1X transfer buffer (25mM Tris base, 190mM Glycine, 20% MeOH, 0.05% SDS), and transferred (Bio-Rad) at 100V for 90 min. Membranes were taken through a standard immunoblot protocol followed by enhanced chemiluminescent detection (Crescendo ECL, Millipore) using a Lumimager (Fuji, LAS-3000). Primary antibodies: α-tubulin (1/1000, Sigma), α-flag (1/1000, Cell Signaling Technologies), α-myc tag (1/1000, Cell Signaling Technologies), GFP (1/2500, Santa Cruz), β-actin (1/1000), α-PAR (1/1000, Trevigen) and α-Axin (Feng et al., 2014; dT20 *Drosophila* Axin antibody, 1/250, Santa Cruz). Secondary antibodies: anti-mouse, anti-rabbit, anti-guinea pig (1/5000, Jackson Labs).

### Immunofluorescence

Adult intestines were dissected in 1xPBS and fixed in 4% paraformaldehyde in PBS for 45 min at room temperature. Tissues then were first washed with 0.1% Tween 20-PBS, second washed with 0.1% TritonX-100-PBS and finally permeabilized in 0.5% TritonX 100-PBS for 30 min. Following the blocking with 10% BSA in 0.1% Tween 20-PBS for 1h at room temperature, primary antibody incubation in 10% BSA in 0.1% Tween 20-PBS was performed for overnight at 4°C. 3X 5 min 0.1% Tween 20-PBS washed intestines then were incubated in secondary antibodies for 1h at room temperature. Specimens were finally mounted in Fluoromount-G (Southern Biotech) and analyzed with confocal imaging. Primary antibodies: mouse anti-Arm (Wang et al., 2016; N2 7A1, DSHB, 1:50), mouse anti-Prospero (Wang et al., 2016; MR1A, DSHB, 1/50), mouse anti-GFP (GFP-12A6, DSHB, 1/100), mouse anti-β-galactosidase (Tian et al., 2016); 401A, DSHB, 1/100), mouse anti-Delta (Wang et al., 2016; C594.9B, DSHB, 1/100), rabbit anti-phosho-S10-Histone3 (Wang et al., 2016; 06-570, Millipore, 1/1000). The secondary antibodies were Alexa fluorophores (Thermo Fisher Scientific) and diluted as 1/1000.

### Quantification of Stat-GFP immunostaining intensity

Images from R5 region were taken with a 63x objective. Each STAT-GFP^+^ stem cell was identified using Imaris software (Bitplane). The main intensity in those cells within a field (40 μm x 40 μm) surrounding an *Iduna* mutant clone or an equal field at least 50 μm away from the mutant clone was measured. The relative intensity was calculated and shown in the figure (Wang et al., 2016). Statistical analysis was performed with Prism software (GraphPad).

### Immunoprecipitation

S2R+ cells were seeded at 5x10^6^ cells/10 cm^2^ culture plates and incubated overnight at 25°C. Cells were then co-transfected with 5µg of each plasmid by using Mirus-insect transfection reagent. Negative control was transfected with empty plasmids. 48 hours later, transfected cells were harvested. The cell pellets were washed within cold 1X PBS. This step was repeated 3 times. Pellets were re-suspended in 600µl 1% Triton X-100 lysing buffer. Re-suspended pellets were incubated on ice for 15 min and mixed gently and periodically. Total lysates were centrifuged at 13,000 rpm at 4°C for 30 minutes. The supernatant was removed and 100µl was stored as total lysate. 25µl Protein A/G (Thermo Scientific) beats were washed with lysing buffer for 3 times. 200µl supernatant was incubated with the mixture of washed protein A-G beads on a rotator at cold room for 30 minutes. In a parallel way, 25µl Protein A/G was washed with lysing buffer for 3 times. At the end of incubation period, beads-supernatant mixture was centrifuged at 2,000 rpm at 4°C for 1 minute. Pre-cleaned supernatant was collected and added to beads. Antibody was added to supernatant-beads and incubated at cold room on a rotator for 4 h. Beads-supernatant-antibody mixture was centrifuged at 2,000 rpm at 4°C for 1 minute and beads were washed with lysing buffer for 3 times. In the final step, beads were re-suspended in 50µl of 3X sample buffer to perform immunoblotting.

### Recombinant protein purification from S2R+ cells

S2R+ cells were seeded at 5x10^6^ cells/10 cm^2^ culture plates and incubated overnight at 25°C. Then, flag or myc-tagged gene of interests were transfected and based on the small tag, a recombinant protein was immunoprecipitated with flag or myc agarose beads as described above. Finally, by using flag or myc peptides, tagged proteins were eluted and quantified by BCA (Pierce, ThermoFisher Scientific).

### Quantification and statistics

ISC quantification, dissected midguts were stained with Armadillo and Prospero. Images of the R5 region (Buchon et al., 2013) were obtained with a 63X objective and total number of Arm^+^/Pros^-^ cells in a field were counted. Quantifications of immunoblot were done with Image J. Student *t-test* and ANOVA were used as statistical analysis and those were done with Prism (GraphPad) software.

## Author Contributions

HS and YG designed the concept of the study. YG performed all experiments. YG and HS jointly wrote the manuscript.

## Acknowledgments

We would like to thank all previous and current members of the Steller Lab for their helpful suggestions and discussions, especially Adi Minis and Junko Shimazu for critical reading of the ms. We also thank Drs. Norbert Perrimon, Jean-Paul Vincent, Wei Du, Jessica Treisman, and Steven X. Hou for sharing their published *Drosophila* lines, the Bloomington Stock Center and the Vienna *Drosophila* Research Center for the fly stocks, and the *Drosophila* Genomics Resource Center and Developmental Studies Hybridoma Bank (DSHB) for reagents. This work was supported by NIH grant RO1GM60124 to H.S.

The authors declare no competing interests.

## Supplementary Figure Legends

**Figure S1:**
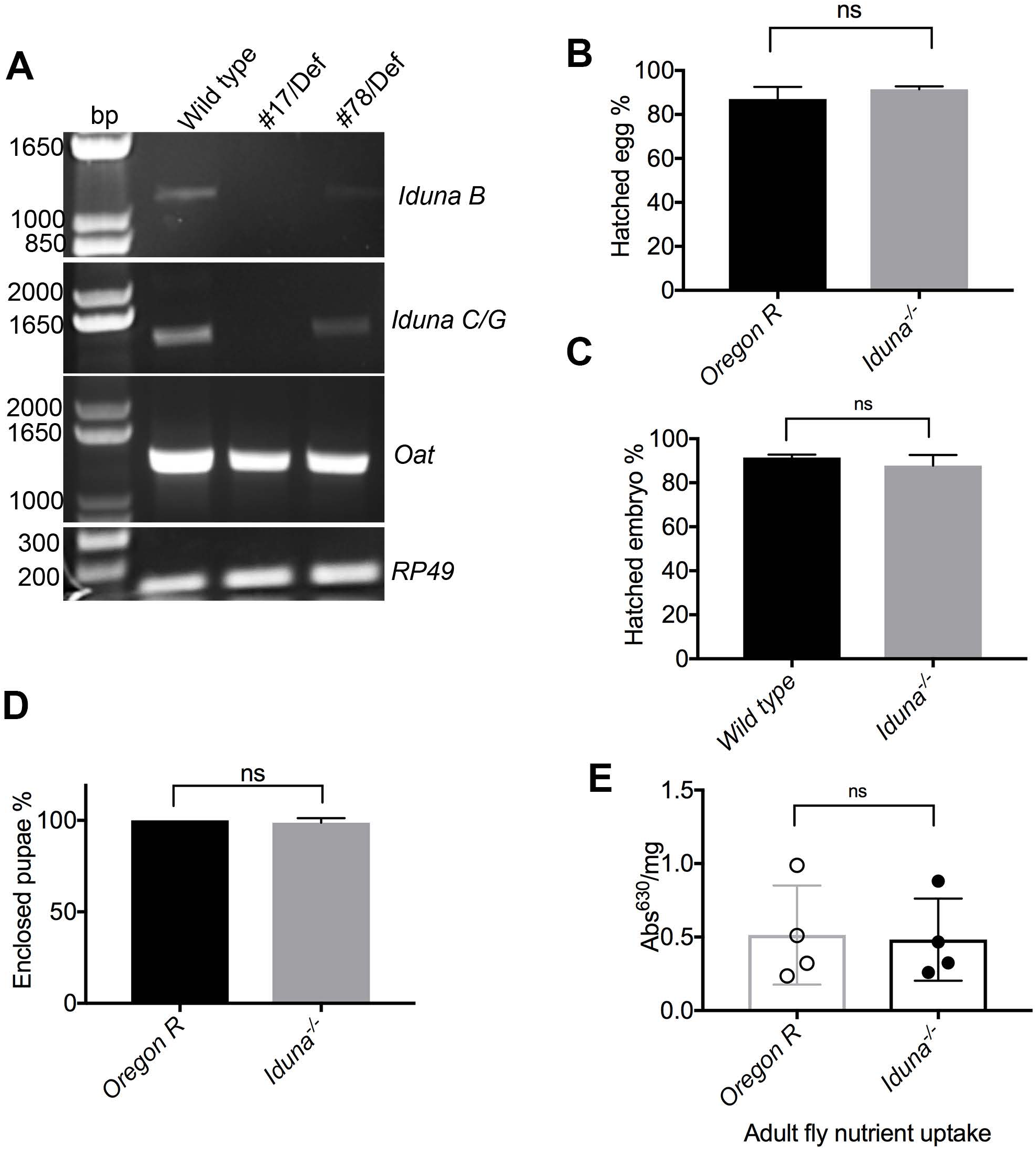
*Iduna* inactivation does not cause developmental defects in *Drosophila.* **A-** Identification of *Iduna*^*17*^ and *Iduna*^*78*^ mutant alleles. *Iduna*^*17*^ had no detectable *Iduna* transcripts, and Iduna^78^ had severely reduced *Iduna* mRNA based on RT-PCR. 7 day-old adult females were analyzed for expressions of *Iduna, Ribosomal protein 49* (a house keeping gene) and *Ornitate aminotransferase.* **B-** *Iduna* mutants did not have defects in hatching their eggs.n>250 for each genotype. **C-** *Iduna* transheterozygous mutants did not have defects in hatching their eggs. n>100 for each genotype. **D-** *Iduna* mutants do not have nutrient uptake *Iduna* mutant larvae could be pupated and enclosed to adult *Drosophila.* n>100 for each genotype. **E-** There is no decrease in food intake in *Iduna* mutant when flies kept under the regular diet. Quantification of adult fly nutrient uptake by a calorimetric assay from regular dietary condition in *Iduna* mutants and Oregon R.

**Figure S2:**
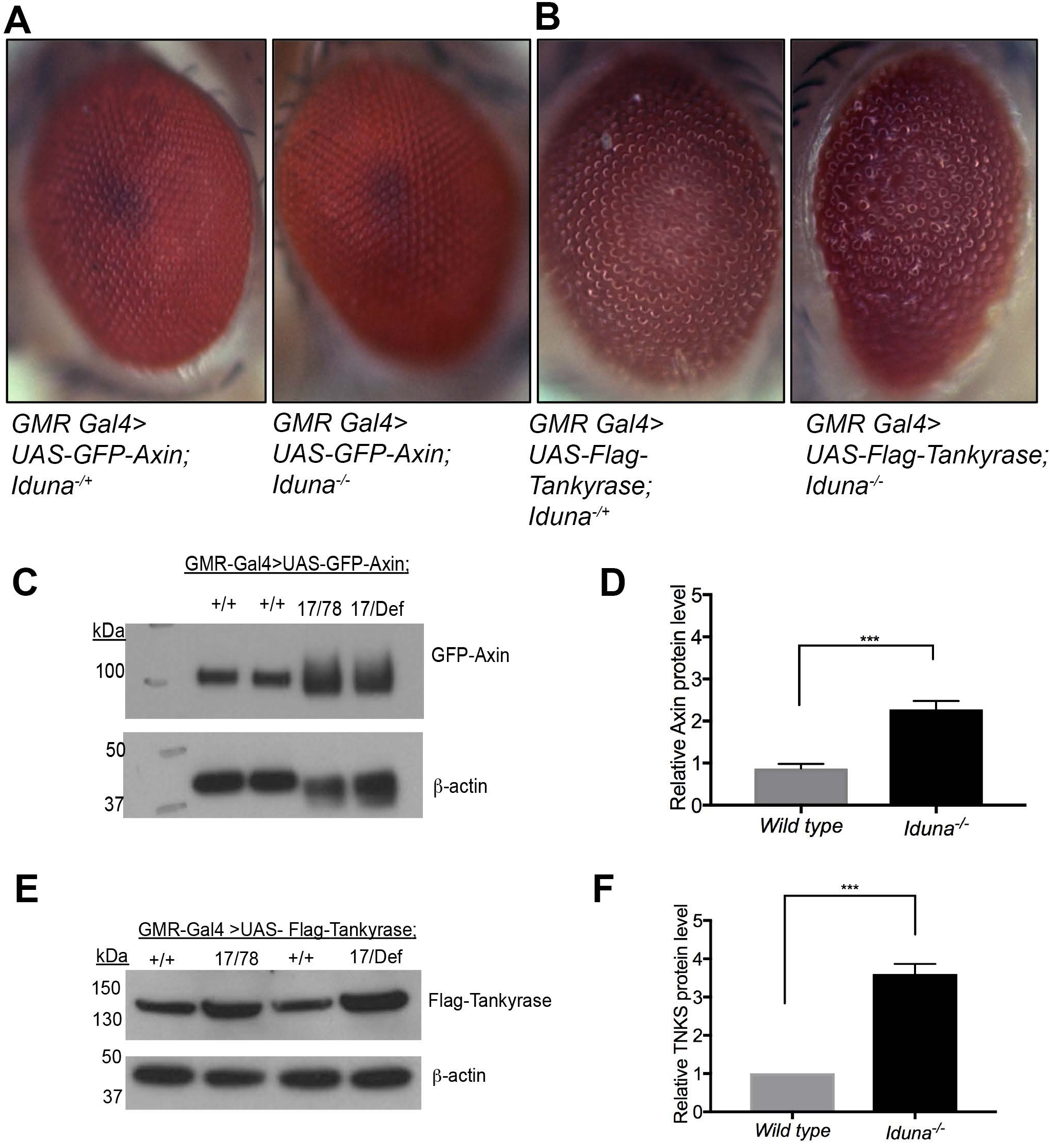
*Iduna* depletion leads to increased Axin and Tankyrase protein levels. **A-** Mis**-** expression of GFP-Axin in adult eyes did not result in obvious eye phenotype. Five day-old adult male eyes were imaged. **B-** Mis-expressed Tankyrase however led to rough eye. *Iduna* inactivation promotes the eye phenotype. Five day-old adult male eyes were imaged. **C-** *Iduna* inactivation leads to mis-expressed GFP-Axin elevation. A UAS-GFP-Axin reporter transgene was expressed under *GMR* driver in the *Iduna* mutants or wild type. **D-** *Iduna* mutants had 2.5-fold increased GFP-Axin protein. Quantification of mis-expressed GFP-Axin immunoblottings. Results are based on two repeats of independent replicates and Axin protein levels were normalized to β-actin. **E-** Flag-tagged mis-expressed TNKS protein accumulates in *Iduna* mutants. A UAS-Flag-TNKS reporter transgene was expressed under the *GMR-Gal4* driver in *Iduna* mutants or wild type. **F***-Iduna* mutants had 3.5-fold increased levels of flag-tagged TNKS. Quantification of Flag-TNKS immunoblottings. Five day-old adult male heads were dissected and 20µg of total protein lysates were analyzed by immunoblotting to assess the levels of GFP-Axin or Flag-tagged TNKS using α-GFP or α-flag antibodies, respectively. Western blot quantification was performed based on two independent experimental replicates, and protein levels were normalized with β-actin. Oregon R flies were used as a wild type control. Flies were fed with regular diet at 24-25°C.

**Figure S3:**
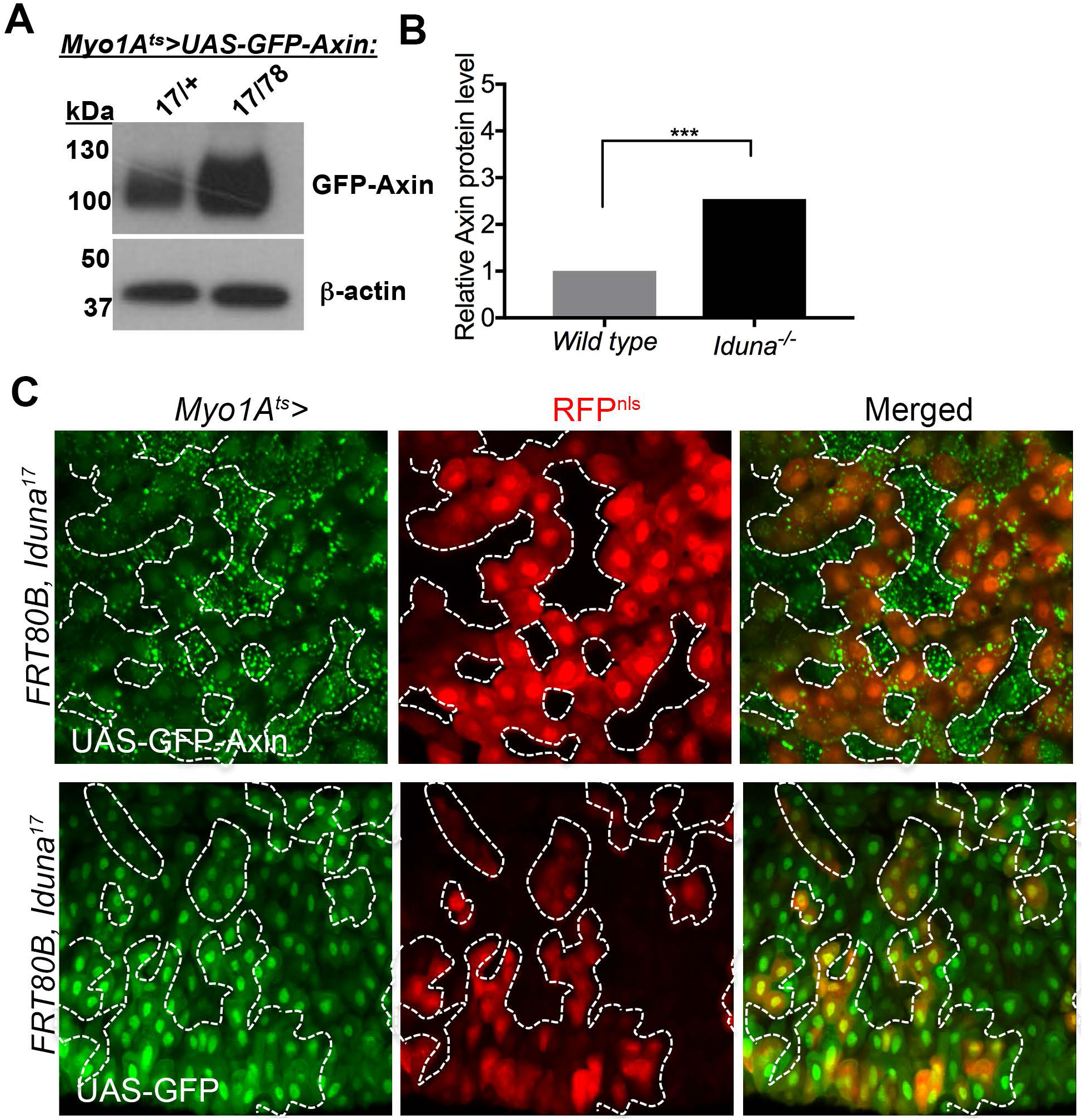
*Iduna* inactivation causes increased mis-expressed Axin protein levels in the midgut. **A-** *Iduna* mutant midguts had elevated mis-expressed GFP-Axin protein. Midguts of 7 day-old adult females, which were expressing GFP-Axin under the temperature sensitive *Myo1A-Gal4* driver, were dissected, lysed and analyzed by GFP immunoblotting. *Iduna* mutants had more Axin protein compared to the wild type. 20µg total intestinal lysates were analyzed by GFP immunoblotting and α-tubulin was used as a loading control. **B-** Loss-of-*Iduna* resulted in 2.2-fold GFP-Axin accumulation in the midgut. Western blot quantification was performed based on two independent experimental replicates, and protein levels were normalized to α-tubulin. **C-** *Iduna* mutant clones have elevated mis-expressed GFP-Axin compared to their WT neighbours. A UAS-GFP-Axin transgene was expressed under the temperature sensitive *Myo1A-Gal4* driver in the FRT80B-*Iduna*^*17*^ mutant. Midgut mutant clones were induced during larval development by daily incubation at 37°C for 1h. Adult female FRT80B-Nls-Red/FRT80B-*Iduna*^*17*^ flies were collected after eclosion, incubated at 29°C and analyzed on day 7. Unlabeled cells represent *Iduna* mutant clones, whereas cells stained for nuclear RFP are either wild type or *Iduna* heterozygous. *p<0.001* is indicated as *** and *p<0.0001* was marked as ****.

**Figure S4:**
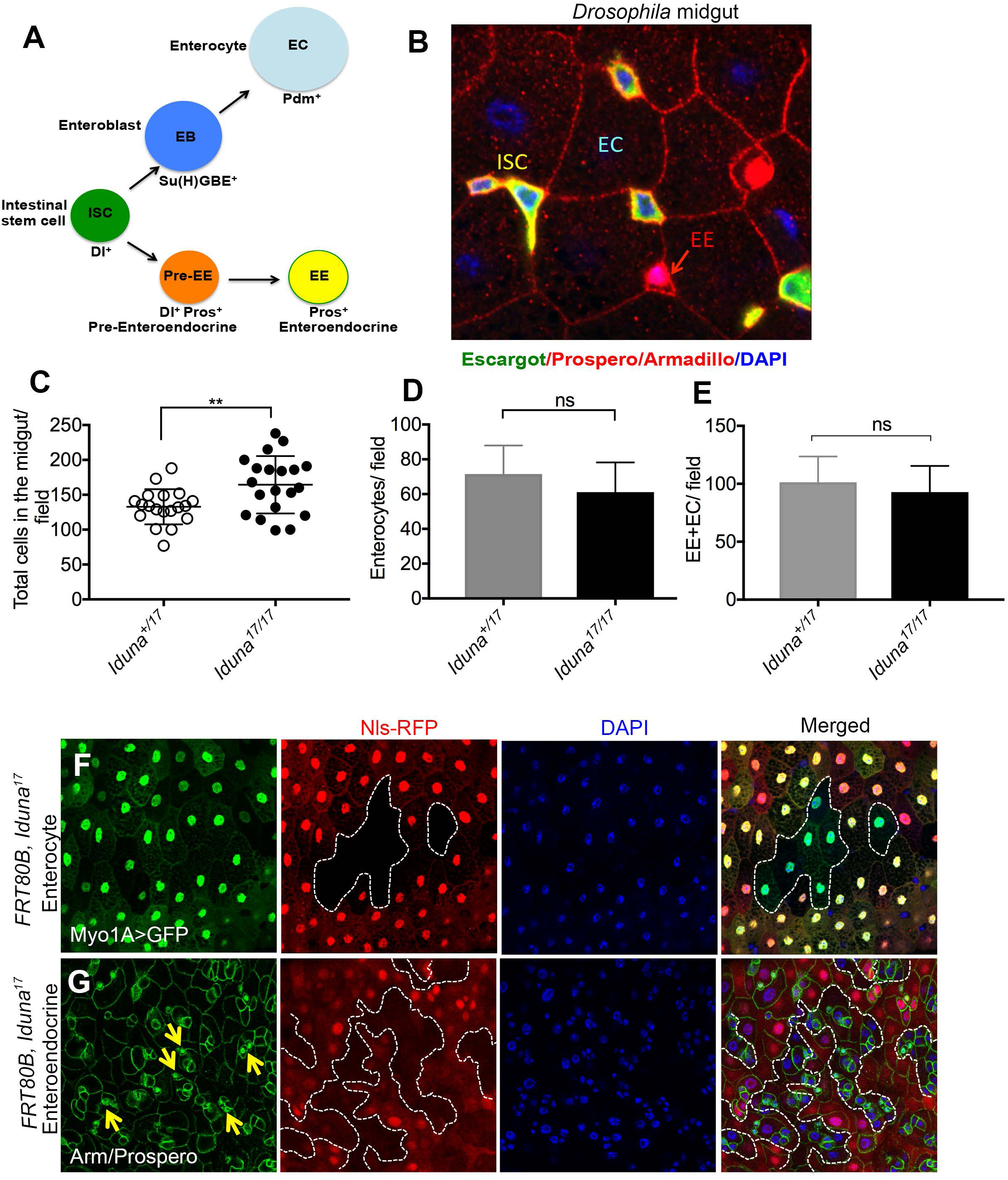
Iduna is not required for differentiation of ISC. **A-** Intestinal stem cells in the *Drosophila* midgut give rise to enterocytes and enteroendocrine cells. ISCs give rise to two different types of daughter progenitors, undifferentiated enteroblasts (EBs) and pre-enteroendocrine cells (Pre-EE). EBs and pre-EEs differentiate into enterocytes (ECs) and enteroendocrine (EEs) cells, respectively. **B-** ISCs and ECs can be distinguished by their cell sizes, high level of membrane-associated Armadillo, and lack of nuclear Prospero. On the other hand, differentiated EEs are small and can be identified by nuclear Prospero staining. Finally, small-sized ISCs are stained with Armadillo but not with nuclear Prospero. In the image, small ISCs were co-stained with escargot (green), and Armadillo (red). On the other hand, small EEs were shown with the red arrow and stained with Armadillo (red) and nuclear Prospero (red). Bigger cells are enterocytes whose cell membrane is stained with Armadillo (red). DAPI staining in blue marks the cell nucleus. Flies were fed with regular diet at 24-25°C.*Iduna* mutants contain slightly more cells in their midgut. It appears that an increase in ISC number is responsible for this effect. **C-E-** No differences in the numbers of enterocytes and enteroendocrine cells were observed in *Iduna* mutants. **F-** Iduna is not required for the differentiation of stem cells into enterocytes. Enterocytes were marked with GFP expressed under the control of enterocyte specific driver, *Myo1A-Gal4*. 7d-old mutant clones were analyzed. Control cells are shown with nuclear RFP, and *Iduna* mutant clones are marked with a white line. **G-** Genetic depletion of *Iduna* did not affect the differentiation of ISCs into enteroendocrine cells. Mutant clones in 7d-old adult midguts were analyzed with Prospero staining. Small nuclear Prospero-positive EE cells are labeled with yellow arrows in *Iduna* mutant clones. Control cells are labeled with nuclear RFP, and mutant clones are marked with a white line. Flies were fed with regular diet at 24-25°C.

**Figure S5:**
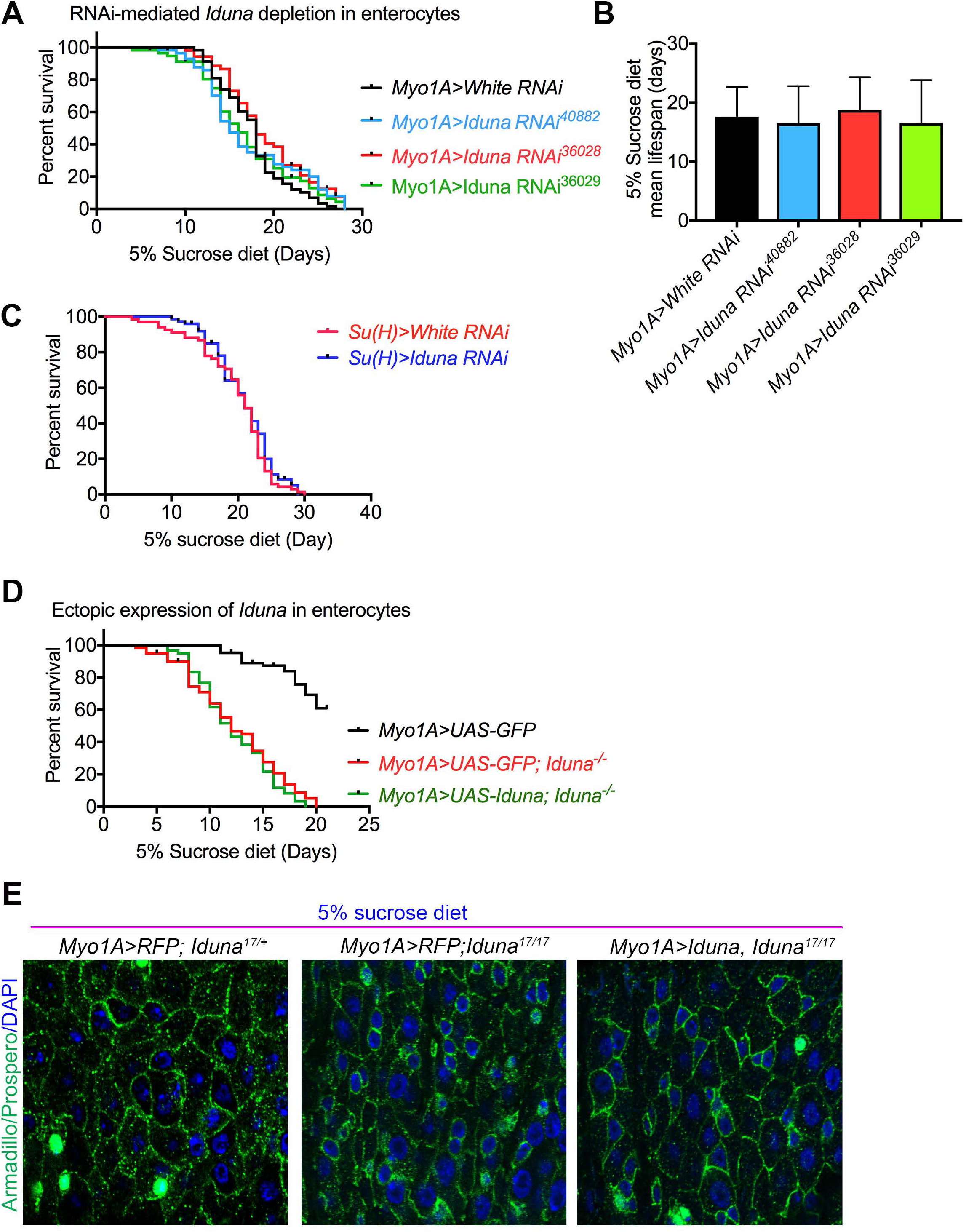
RNAi-mediated *Iduna* depletion in enterocytes does not cause mortality upon nutrient deprivation. **A-** Three different *Iduna* RNAi lines were expressed under the *Myo1A* driver. *White* RNAi was used as a control. **B-** There was no significant change on the mean lifespan between *white* and *Iduna* RNAi expressing flies. n=60 for each genotype. **C-** *Iduna* inactivation in enteroblast does not lead to increased lethality upon 5% sucrose diet at 28°C. n=70 for each genotype. **D-** Ectopic expression of *Iduna* under Myo1A driver in enterocytes does not rescue elevated mortality of *Iduna* mutants under reduced nutrient diet. Two-day old mutant or wild type female flies were collected and kept on 5% sucrose diet at 28°C for the experiment in A-D**. E-** Enterocyte specific ectopic expression of I*duna* rescues the hyper-proliferation of midgut stem cells upon nutrient deprivation. 2 day-old females were collected at 24-25°C and their regular diet was replaced with 5% sucrose diet at 28°C. 9 day-old female flies were examined with α-Armadillo and Prospero antibodies.

**Figure S6:**
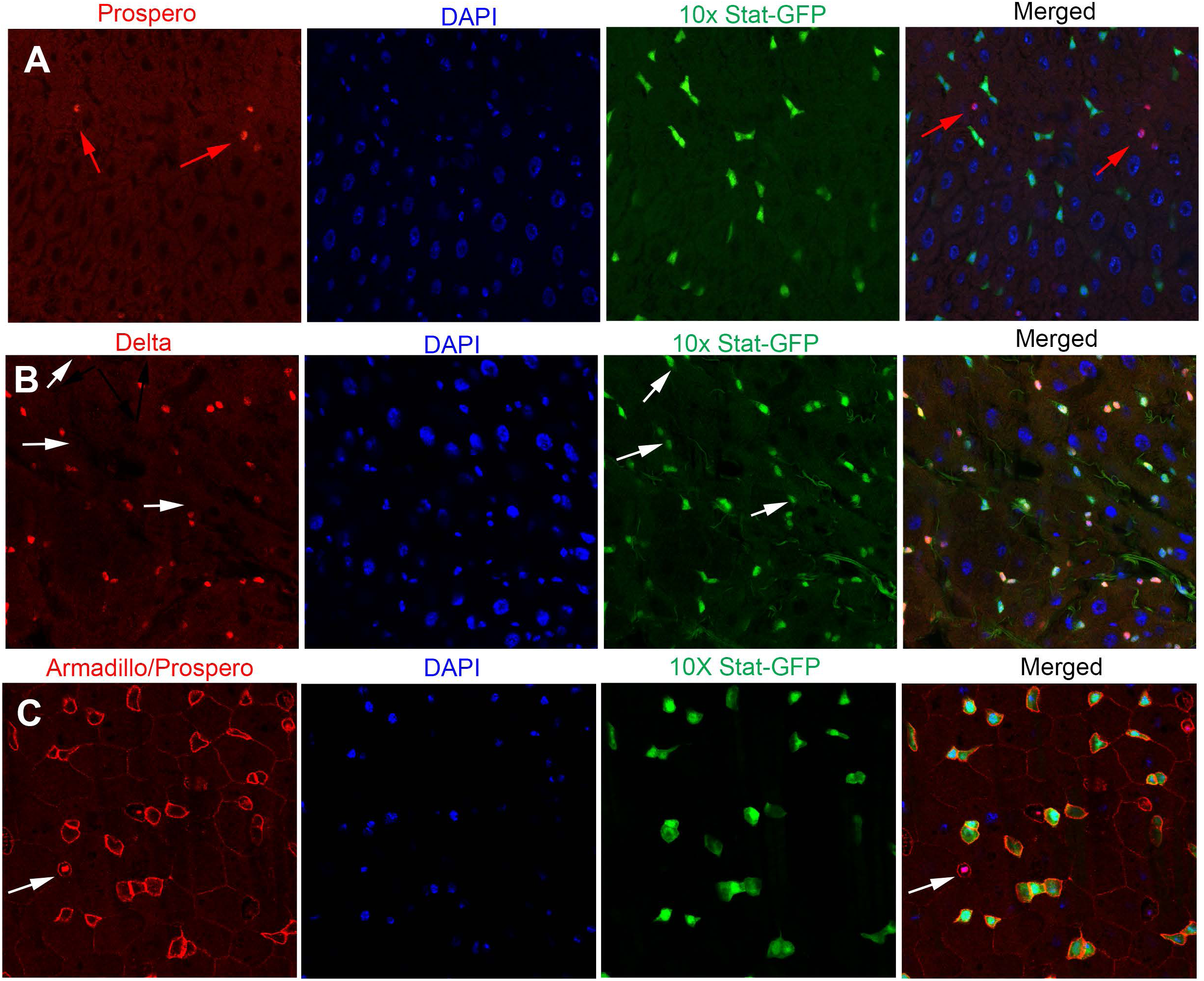
Under regular physiological conditions, the Stat-GFP reporter is mainly expressed in midgut stem cells and enteroblasts. **A-** 10x Stat-GFP reporter was mainly positive in small sized cell populations but Prospero stained enteroendocrine cells were negative for Stat-GFP in the midguts. **B-** Delta-lacZ positive but Prospero negative cells were mainly positive for Stat-GFP expression. It was also indicated with the white arrows that a small population of cells, which were small-sized and GFP positive but Delta-lacZ negative. Those could be undifferentiated progenitors like enteroblast cells. **C-** Armadillo positive Prospero negative small sized stem and progenitor cells have Stat-GFP reporter activity. 5 day-old female flies were dissected and stained with α-Armadillo and Prospero antibodies.

**Figure S7:**
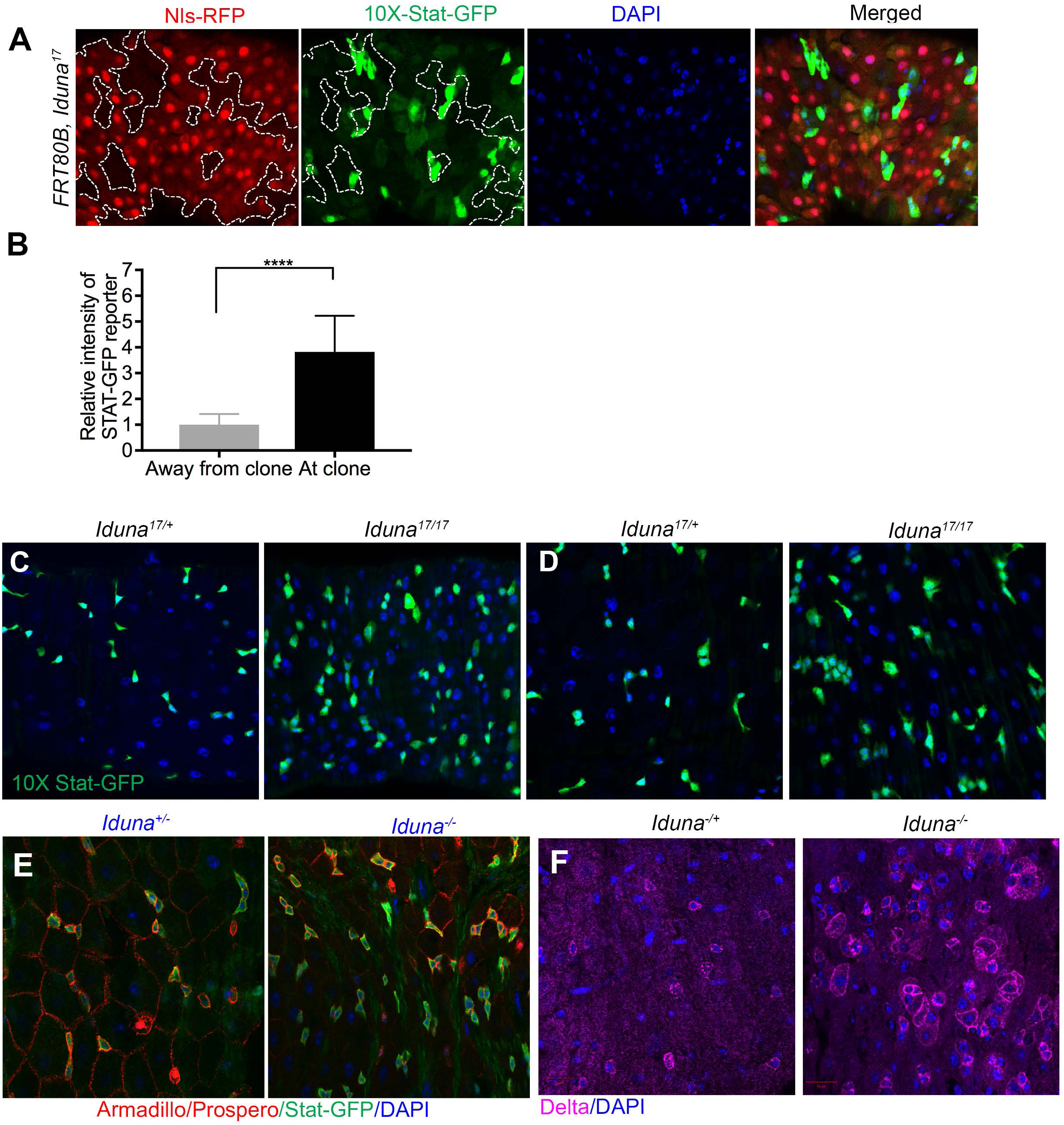
*Iduna* mutants have increased numbers of Stat-GFP positive cells. **A**-*Iduna* mutant clones had elevated Stat-GFP signaling in ISCs. *10X-Stat-GFP* is a reporter for STAT signaling activity. *Iduna* mutant clones displayed strongly increased staining of this reporter. 7 day-old females were dissected and the posterior midguts were analyzed. **B**-Quantification of Stat-GFP reporter expression in *Iduna* mutant clones. GFP intensity was measured with Image J and normalized with control cells. *Iduna* mutant clones showed 4-5 fold higher reporter expression. **C-E-** *Iduna* mutants have more Stat-GFP positive stem cells and progenitors in the midguts. 7 day-old females were dissected and the posterior midguts were analyzed by confocal microscopy. n>6 for each genotype. *p<0.0001* is indicated as ****. **F-** Delta protein is elevated in the midguts of *Iduna* mutants. 7 day-old females were dissected and the midguts were stained for Delta. Flies were collected and kept on regular diet at 24-25°C.

**Figure S8:**
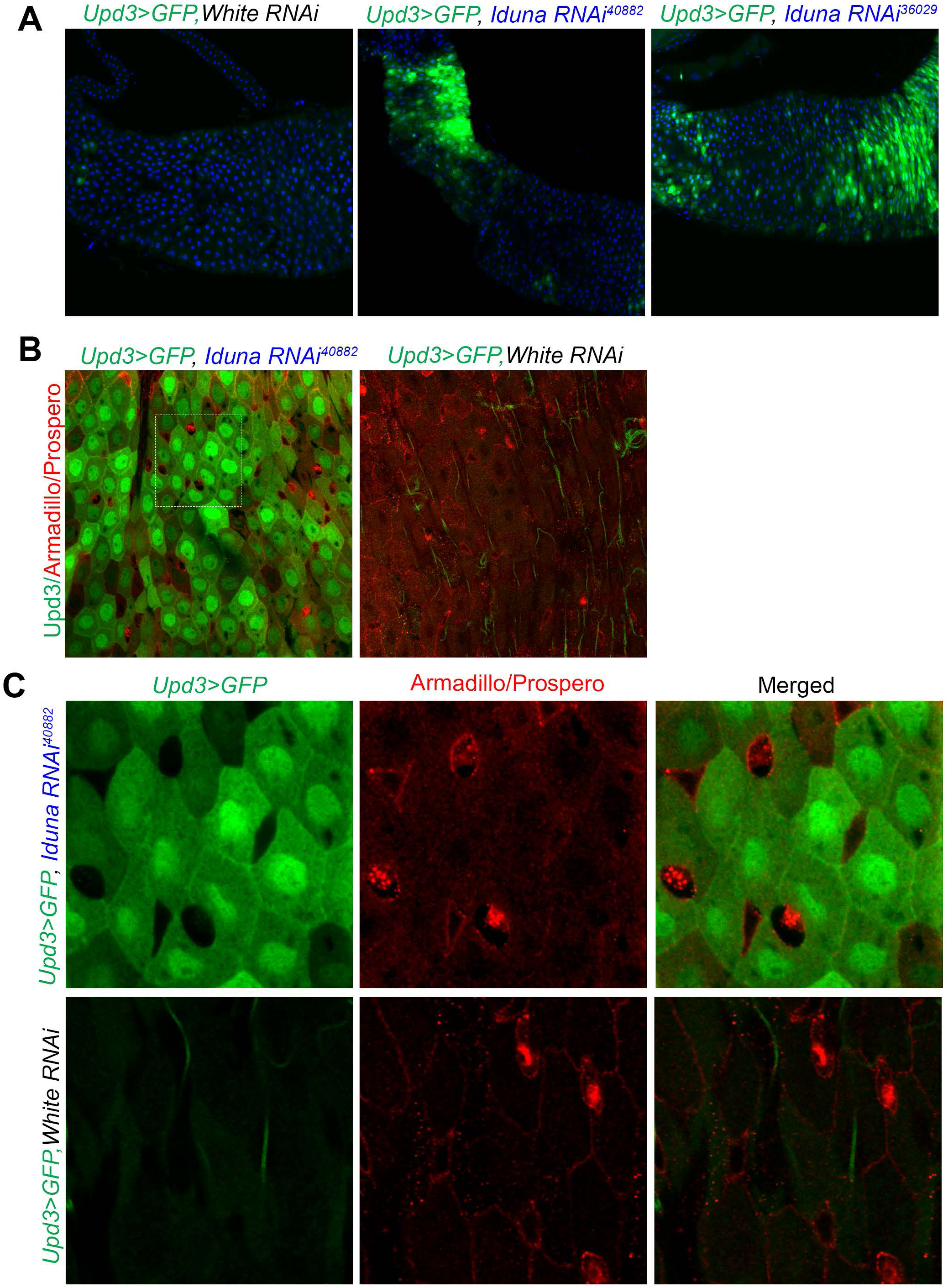
RNAi-mediated *Iduna* depletion increases *Upd3>GFP* reporter activity. **A-** Knock-down of *Iduna* results in upregulation of *Upd3*. *Iduna* was down-regulated by two different UAS-RNAi lines (#36029 and #40882) under *Upd3-Gal4 driver*. *Upd3>GFP* was used as a reporter for *Upd3* gene expression. *White RNAi* served as control. **B-C-** *Upd3 was* upregulated in the enterocytes. Dissected midguts were stained with α-Armadillo and Prospero antibodies. ECs were *Upd3>GFP* expressing cells but EEs and ISCs not. 3 day-old female flies were dissected and their posterior midguts were analyzed by confocal microscopy.

